# Statistical Molecular Interaction Fields: A Fast and Informative Tool for Characterizing RNA and Protein Binding Pockets

**DOI:** 10.1101/2025.04.16.649117

**Authors:** Diego Barquero Morera, Giovanni Mattiotti, Alexander Kocev, Amshuman Rousselot, Louis Meuret, Lucas Rouaud, Hubert Santuz, Marc Baaden, Antoine Taly, Samuela Pasquali

## Abstract

Developing a physical understanding of the interactions between a macro-molecular target and its ligands is a crucial step in structure-based drug design. Although many tools exist to characterize protein-binding pockets in silico, this is not yet the case for RNA, which has only recently been recognized as a suitable target for small ligands. Molecular Interaction Fields (MIF) are a useful tool to characterize the interactions of a given binding pocket. However, classical MIFs heavily rely on the use of probes, which makes their calculation accurate but very specific to the binding partners in question. We develop here a simple version of MIF, that we call Statistical Molecular Interaction Fields (SMIF), based on functional forms inspired by coarse-grained models and parametrized based on PDB structures and previous statistical analysis of the main form of interactions typical of macromolecules, namely hydrogen bonding, stacking, and hydrophobic interactions. We show that these fields, despite their simplicity, are very informative and overall in agreement with pharmacophoric models. Thanks to a carefully optimized code, our calculations are fast and can be performed in bulk on a large set of binding pockets or even on a full macromolecule. As shown in a few representative examples, the latter possibility opens the way to the analysis of systems as large as 20000 to 80000 atoms in relation to the surrounding environment, i.e., a lipidic membrane, a small ligand, or another macromolecular partner, allowing for a detailed visualization of the possible interactions. The complete software and its documentation are available here: https://smiffer.mol3d.tech/

## 1. Introduction

Small molecules modulating the behavior of biomolecular targets are the current main form of medication for most diseases and, although other forms of drugs exist such as biologics and therapeutic peptides, small-molecule drug design is still a very active research field [1, 2]. To this day, almost all approved drugs target proteins such as enzymes, receptors, ion channels, and more, but with the discovery of the functional role of RNA in many cellular processes, RNA molecules (non-coding RNA and mRNA) have been identified as possible targets [3] and research to design RNA drugs is growing in importance [4, 5, 6].

The open challenges in drug design are to improve the efficacy of a drug by finding new compounds with high affinities and specificity, and accounting for the flexibility of the target [7]. From a computational point of view, this problem is being tackled both using traditional molecular modeling approaches based on docking, scoring functions refinement, and molecular dynamics simulations[8, 9], and, more recently with the help of artificial intelligence [10, 11].

The development of drugs specifically targeting RNAs has recently opened a whole new research field. Up until recently only antibiotics were known to target RNA (in the ribosome), but it is now clear that non-coding and mRNAs constitute a new pool of possible targets, complementary and possibly wider, than the space of druggable proteins. Only roughly 2% of the human genome codes for proteins, while about 70% translates into ncRNA and most viruses threatening new pandemics have a single stranded RNA genome [12]. Drugs targeting RNAs must have their own specific properties, different from those targeting proteins. This requirement arises from the nature of RNA that are highly charged molecules constituted of nucleotide building blocks with less structural variability than amino-acids. The smaller RNA alphabet and the similar physico-chemical nature of the four nucleotides raise the problem of specificity of a drug in binding to a target [3]. Moreover, RNA are often subject to post transcriptional modifications such as methylation [13], which can substantially alter their structure and binding modes, and whose sites can become targets for small ligands and protein interactions [14, 15, 16]. In general, the characteristics of RNA binding sites are less understood than their protein counterparts [17, 18], but the tools developed to study in silico the interactions between macromolecules and small ligands were mainly developed for proteins and later adapted for RNA [19].

In order to gather a rational understanding of these interactions, of the role of flexibility, and of the specific differences between protein and RNA drugs, physical modeling is necessary and provides a guiding light to develop advanced computational methods, including AI-based ones. As RNA structural data is still rather limited to be exploited for machine learning, the rules of the physical interactions of these molecules should be generally valid and transferable.

In this work, we present a novel approach for looking at molecular interactions based on the concept of interaction fields computed with simple functional forms based on the statistical analysis of existing complexes. Our approach has the advantage of allowing fast calculations and insightful visualizations. It has the potential to become a unique investigative tool integrated with molecular visualization software, enabling researchers to intuitively gain insights into the systems under study and design more targeted drug development strategies.

*Molecular Fields.* A typical approach used in molecular simulations and in structural analysis is to consider pairwise interactions between selected elements in the system to help rationalize the binding modes of a receptor and a ligand or between two biomolecules. An alternative point of view, more physics-inspired, is to focus on just one object, such as a macromolecule, and to consider the interaction “fields” it generates and that dictate how it might interact with other partners. A field is an intrinsic property of the object and it consists in the modification of the properties of the space around it, induced by the presence of the object itself. This is the case for example of the gravitational field of the Earth, and of the electrostatic field generated by a point charge.

The concept of an object’s field can be applied to molecular interactions focusing only on one system and considering the possible interactions it can generate, independently of a specific binding partner. This approach is particularly interesting in the context of a biomolecule to be targeted with a small ligand: if we can characterize the properties of a binding pocket, we can then make hypotheses on the type of ligand that can bind to it and how. The concept of Molecular Interaction Fields (MIF) was introduced already in the ’70s and implemented initially in the ’90s [20]. Molecular fields are typically obtained using probes that are positioned on a grid sampling the entire space of interest and are given as the interaction energy between the probe and the receptor computed from a molecular force field [21, 22]. The latest implementation of the GRID software, in 2021, accounts for over 70 different probes [23]. Over the years, molecular fields have been applied extensively in both ligand-based applications to assess ligand similarity, virtual screening and ADME predictions, and in structure-based applications to characterize and detect binding pockets, and for docking [24]. The widely used Autodock software computes atomistic force field MIFs of a binding pocket with the atom types of the ligands as probes and then uses the fields to guide docking poses [25]. With a similar approach, higher resolution quantum calculations were used to characterize non-bonded interactions with high spatial precision, focusing on the interactions of small subsets of atoms [26]. All these approaches to compute MIFs are based on the use of probes, making their calculations energetically consistent, and the information obtained specific to the complex, as it depends on the nature of two partners. Because of the high accuracy, these calculations can be rather expensive and they are typically performed only in small regions of the molecule, such as a binding pocket.

Wanting to develop a fast and general method to build an intuitive understanding of possible molecular interactions, we have defined simplified molecular interaction fields, based on a statistical analysis of the structural database of proteins and nucleic acids, and that take inspiration on coarsegrained molecular models [27, 28]. Speed of these calculations is crucial if we want to apply them fluidly in interactive visualization and eventually integrate these calculations into common visualization software. Speed is particularly relevant to study RNA molecules. Given their inherent flexibility, the calculations may need to be performed on an ensemble of structures and not just a single one, to bear relevance.

Over the years we worked extensively on the development of visualization software with the specific aim to allow the user to build an intuition on the system under study [29, 30]. Our UnityMol software was developed with the ability of performing interactive simulations and allows an immersive approach with virtual and augmented reality [31, 32, 33]. Combining such virtual reality approaches and SMIF visualization could be a powerful new tool for drug design [34].

For our purposes, field calculations don’t need to be as detailed as those evaluated from atomistic force fields. We therefore define effective statistical potentials (that we name SMIF, for Statistical Molecular Interaction Fields) for hydrogen bonding, stacking and hydrophobic effect. We rely on the APBS (Adaptive Poisson-Boltzmann Solver) software for calculations of electrostatic fields generated by point charges [35]. Two important differences between our fields and the original MIFs are that we do not use probes, making our calculations particularly efficient and our results general, and we adopt a coarse view, which in particular allows us to easily define a stacking term. A comparison between original MIFs and our SMIFs on two systems (one protein and one RNA) is presented in Supplementary Information.

In this manuscript, we show that, for known complexes of a biomolecule and a small ligand, the statistical fields computed solely on the knowledge of the binding pocket are in excellent agreement with the positions of the pharmacophores obtained for the ligand. This observation indicates that even fields built with the simple approach that we propose are highly informative of the interactions accessible to the biomolecule. It is important to notice that our excellent results hold for both proteins and RNAs.

Since calculations are inexpensive and the correspondence between computed and known properties of the biomolecule is satisfactory, we can use SMIFs to investigate beyond the spatial boundaries of a given binding pocket and extend the analysis to the whole macromolecular systems. This extension can have useful applications in the analysis of interfaces of an entire biological macromolecule and in the search of possible binding pockets themselves. As we show in an example, in the context of the open problem of RNA drug design, exploiting the information from SMIFs could be a valid approach to identify RNA binding pockets. Our approach can be used to rationalize the interaction between large macromolecules, as we show in an example of a ribosomal subunit interaction with an Internal Ribosomal Entry Site.

In this work, we first present how SMIFs are defined and built. We then present a few examples where we visualize the fields, followed by the benchmark with pharmacophores. We show how comparing the fields of pockets to the fields around the whole macromolecule can help characterize the nature of the interaction that the pocket can accommodate and score RNA binding pockets. We conclude with two examples of use of SMIFs on entire macromolecular systems where we show how the fields can help characterizing interfaces of large molecular complexes. To make our approach easily accessible to the community, we provide the documented source code, executable builds and a docker container with a set of benchmarks and a series of scripts for visualization. In virtual reality, the fields can be visually explored or used as guidance for interactive docking, using the UnityMol software and provided examples.

## 2. Fields models

Statistical fields are defined to estimate the tendency of single atoms/groups to participate in an interaction. They are built with a top-down approach, which consists in writing a plausible functional form for the interactions, guided by the knowledge of the atomic positions observed in experimental structures, and by assigning the free parameters based on a statistical analysis of a database of structures, or on more detailed calculations and experimental data providing structural information [36, 37, 38, 39]. The molecular interactions we want to account for are electrostatics, stemming from the presence of net or partial charges, hydrogen bonds and stacking, and the hydrophobic effect, stemming from the rearrangement of water molecules.

From experimental structures and molecular simulations, we know that hydrogen bonding and stacking are present when the atoms adopt well defined geometries. Our potentials are chosen to have minima in the optimal geometry and to rapidly go to zero when atoms are far apart, to obtain well localized interactions. For our fields we choose Gaussian functional forms as they have these properties and are easy to interpret as probabilities.

*Stacking.* The stacking potential field aims at quantifying a property that depends on the position and orientation of the ensemble of atoms composing aromatic rings. The aromatic rings considered correspond to the natural occurrences in residues of proteins (phenyl group for Phe and Tyr; imidazole for His; indole for Trp) and RNA (pyrimidine for Ura and Cyt; purine for Ade and Gua).

For this field we adopt a coarse-grained approach and define a potential based on the location and alignment of a pair of aromatic rings given the distance *r* of their center of mass and their relative vertical position *α* = r · **n**, where **n** is the unitary vector normal to the plane of the ring taken as reference (Fig. 1A). In principle, stacking interactions should depend also on the relative orientations of the planes of the two rings (**n_1_** · **n**_2_), but a statistical analysis of aromatic ring interactions between biomolecules and ligands reveals that there is very little variation from a perfectly parallel orientation. In our model we therefore choose to consider only fully parallel rings and not include this additional variable in the potential. For now the model does not include face-to-edge configuration, but this could be added in a future version after a new statistical analysis of the structures in the PDB.

**Figure 1:**
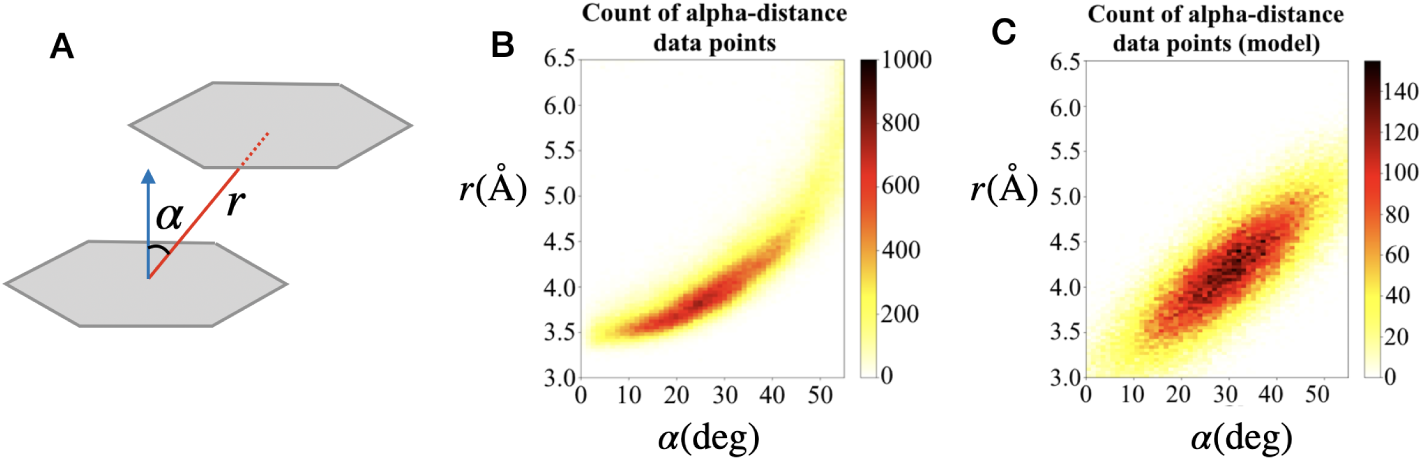
A: model of stacking interactions highlighting the two variables, *r* and *α* used in the definition of the potential field. B: Experimental distribution of values of *r* and *α* extracted from the dataset. C: Modeled potential field for a pair of aromatic rings.

To compute the field, we place a hypothetical aromatic ring centered in each point in space **x**, and we compute its interactions with nearby aromatic rings in the system. The functional form and the parameters of this field were obtained through the analysis of experimentally available structures (details given in Supplementary Information, Section 1). We find that a bivariate Gaussian distribution modeling the interaction of two aromatic rings fits the structural data well (Figure 1:B,C). We therefore define the stacking field as:

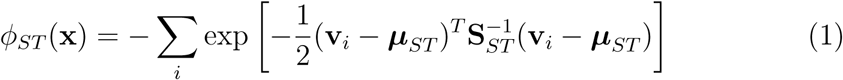

with:

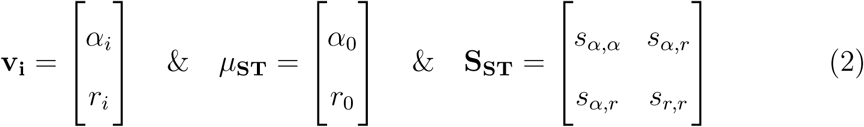

where *i* are the aromatic rings located within 4 standard deviations of the distance (^√^*s_r,r_*) from **x**, *r*_0_ and *α*_0_ are the optimal distance of the center of mass of the rings and the optimal angle for their vertical offset, and the matrix **S_ST_** contains the variances of the Gaussian.

Fitting this function with the distribution obtained from structural data, we determine the model’s optimal parameters to be:

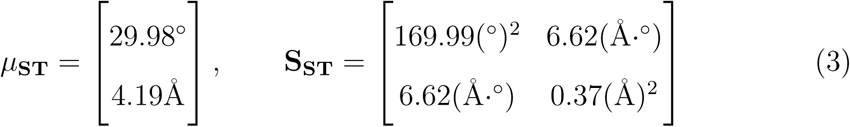

The stacking potential field of a single aromatic group takes the shape of two slightly flattened tori (i.e. “donut” shape), both below and above the aromatic plane (Figure 2:A,B). At each point in space the field is a sum of the potential around each aromatic group, which means that in actual pockets more than one pair of tori can be present. These will naturally overlap with each other in regions where multiple aromatic groups are in close proximity. Points where multiple tori intersect imply a higher propensity to form one or multiple stacking interactions, potentially from both sides of the aromatic plane.

**Figure 2:**
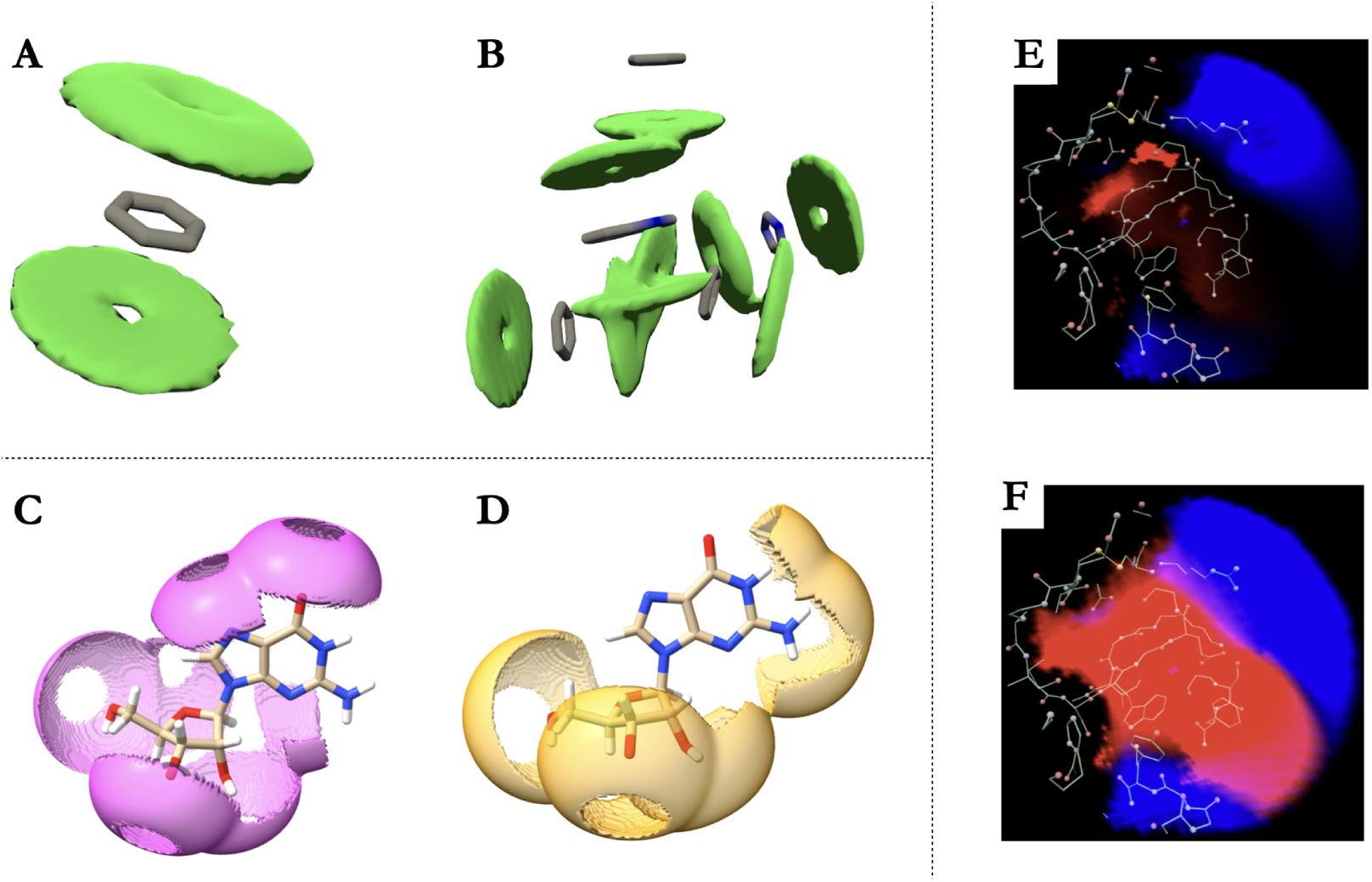
Top left: visualization of stacking fields for a simple aromatic ring (A) and for a binding pocket in which multiple aromatic rings are present (B). Bottom left: visualization of hydrogen bond acceptor field (C) and donor field (D) for a guanine nucleotide. Right: Visualization of the APBS electrostatic potential (E) and of the logarithmic transformation LogAPBS (F). Regions favoring the interactions with positive charges are in red and with negative charges are in blue.

*Hydrogen Bonds.* Hydrogen bond interactions are usually characterized by the distances and angle between acceptor, donor and the hydrogen atom.

Depending on the experiment that solved the structure, PDB files may or may not include hydrogens explicitly. In both instances we define our hydrogen bond fields by considering the typical distances between donors (D) and acceptors (A), and the angles that are formed involving the antecedent atoms bound to the donors (AD) and the antecedent atoms bound to the acceptors (AA).

More specifically, we define two fields: one for the hydrogen bond donors (HBD), and another for the hydrogen bond acceptors (HBA). The HBA field depends on the distance from the acceptor (*r* := |**x** − **x***_i_*|) and the AA A – D angle *β*. The HBD field also depends on the distance from the donor, while the angle *β* it considers depends on whether the structure has hydrogens present or not. When hydrogens are missing, as in structures from X-ray crystallography, the DA D A angle is taken into account. When hydrogens are present, as in structures from NMR experiments, the D H A angle is considered instead.

We define the hydrogen bonding field as the product of two normal distributions of distance and angle as:

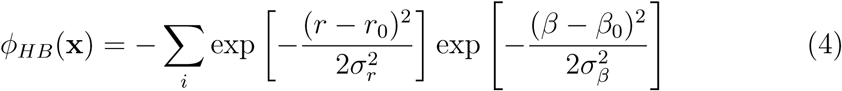

The definition of *β* determines the values of the parameters used in the formula of HBA and HBD. Based on statistical studies on these same angles and distances reported in [37, 38, 39], we take the parameters to be *r*_0_ = 2.93υ and *σ_r_* = 0.21υ. For HBA we take *β*_0_ = 129.0^◦^ and *σ_β_* = 20^◦^. For HBD, when hydrogens are present in the PDB structure we take *β*_0_ = 180^◦^ and *σ_β_* = 30^◦^. When hydrogens are missing there are two possible scenarios depending on whether the bond between the antecedent the missing hydrogen would be. For sp3 aliphatic bonds, such as OH and NH_3_, the donor could rotate around its antecedent, hence the missing hydrogen position is uncertain and only known to be present on a ring around the donor (similarly to what we have for HDA). In this case we take *β*_0_ = 109.0^◦^ and *σ_β_* = 20^◦^.

For bonds whose plane is fixed (such as certain NH_1_ and NH_2_ sp2 bonds) or that are under other geometrical constraints (like the case of NH in an Nterminal proline’s ring), the position of the missing hydrogen can be inferred from the nearby atoms, since the plane on which the bond occurs is known. In these cases, the direction in which the missing hydrogen should be is computed and then the same parameters used for structures with hydrogens are employed.

The SMIFER code detects the presence or absence of hydrogens automatically and prioritizes using them when present. Even if the structure has hydrogens, if any particular donor is missing its hydrogen for some reason, the model falls back to using the alternative method for that atom.

Similarly to the stacking potential, several rings will most likely overlap with each other when studying actual pockets (see Figure 2:C,D).

*Hydrophobicity.* To properly consider hydrophobic effects one would need to dynamically analyze the interactions between the receptor and water. However, for proteins, the propensity of different amino acids toward a hydrophobic or hydrophilic behavior is well known. Indeed, hydrophobicity is usually defined as a property of amino acid residues and visualization software uses hydrophobicity scales to color different amino acids accordingly. A variety of hydrophobicity scales exist and they are often rooted on the idea of transfer energy needed to put a given amino acid in a membrane-like environment, which could be derived from experiment or computation.

Similarly to what is done for hydrogen bonds and stacking, but without the angular dependence, for each atom in the pocket we define a Gaussian field and we assign the height of the Gaussian based on the Wimley and White hydrophobicity scale [40]. This scale assigns an hydrophobicity value *K_i_* to each amino acid; when this is negative, it means that the residue is hydrophilic. Projecting the property of the residue to each atom belonging to it allows preserving the detailed geometry when computing volumetric properties of the pocket. For each point in space we then add the contributions of all Gaussians of neighboring atoms. We calculate two different fields: one called *hydrophobic*, that is the sum of all the Gaussians with *K_i_ >* 0, and one called *hydrophilic* (*K_i_ <* 0). For both, the analytical definition follows:

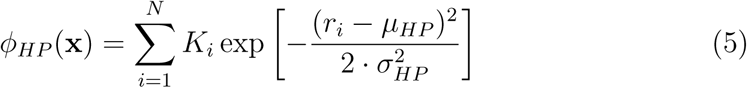

These Gaussians are peaked at a distance *µ_HP_* from atom *i*, equal to 4υ for the hydrophobic field and 3υ for the hydrophilic one. For the first, the choice is inspired by the statistical study that is reported in [36], where the authors show that the typical range of distances between hydrophobic species goes from 3υ up to 7υ, but is peaked around 4υ. For the second, we considered the typical distances between an hydrophilic species and a water molecule (2.5υ 3.5 υ), assuming that the probability of finding a water molecule at a given distance *r_i_* from the hydrophilic species *i* is comparable to the probability of finding another hydrophilic/polar species. Based on these ranges, we also chose *σ_HP_* = 1υ for the hydrophobic field and *σ_HP_* = 0.25υ for the hydrophilic one.

Wimley and White hydrophobicity scale, or similar scales, exist for proteins but not for nucleic acids. Because of the net negative charge of the phosphate group, the backbone of RNA is hydrophilic. The sugar ring is also hydrophilic as it easily forms hydrogen bonds with water because of the OH group. Composed of aromatic rings, bases are hydrophobic and experiments measuring the energy for the transfer of bases from a hydrophobic to a hydrophilic environment give a rough indication of the scale of hydrophobicity between the four bases [41]. From these considerations, we can attempt to give an approximate hydrophobicity scale for the different entities composing a nucleic acid, looking at chemical similarities with amino acids. The negatively charged phosphate group is of similar chemical nature as the side chains of the negatively charged amino acids. We therefore choose the value for the nucleic acid backbone to be the same as that of Glutamic acid, i.e. *K_phos_* = −2.02. The OH group of the sugar resembles the side chain of Serine. We therefore assign to the sugar the same value as Serine, i.e. *K_sug_* = −0.13. Values for the hydrophobic bases are taken to be that of Tryptophan for purines (*K_AG_* = 1.85), since both contain a double aromatic ring, and that of Pheninalanine for pyrimidines (*K_UC_* = 1.13), both containing one aromatic ring. Even though the RNA field computed this way lacks any quantitative precision, this very empirical hydrophobicity scale still enables us to gather an idea of the hydrophobic and hydrophilic regions around these molecules that can be tested against the experimental data of known RNA/ligand complexes (see benchmarks section).

*Electrostatic Potential.* The electrostatic potential field is computed through the Adaptive Poisson-Boltzmann method (APBS), which is a widely used approach to treat electrostatic effects in solution, given an implicit treatment of the solvent [42]. The APBS software generates the electrostatic field on a grid in an assigned volume around the molecule, hence we have simply incorporated the APBS pipeline into our workflow, interpolating the values of APBS field to the grid points defined for our calculations.

Values for the electrostatic field can vary greatly within a system, with very high values occurring in close proximity to atoms with partial charges, while the values quickly dissipate when moving towards the unoccupied volume of the pocket. This wide range of values can be troublesome for visualization. To best exploit information from the electrostatic field, we worked on post processing of the APBS field to generate an output useful for visualization. Two solutions in particular allowed us to improve the visualization of the electrostatic field. The first one, as explained in more details in the next section, consists of trimming regions of space too close to the molecule to be accessible by a ligand. This approach effectively excludes very high values of the field located at the position of the charges. The second, which can at times also be useful, is to convert the field in a logarithmic scale. Since the field can adopt both positive and negative values, this is done by first taking the absolute values of the negative energies and converting both positive and negative values on a logarithmic scale, then applying two empirical cutoffs on the highest and lowest values of both sets (positive and negative), which occur on sparse points, to reduce noise. After subtracting the values of the lower boundary to bring the smallest retained value to zero, we bring back the physical meaning of signed electrostatic potential multiplying by −1 all values that were originally negative. This process generates a second field named LogAPBS. APBS visualization allows to clearly spot maxima and minima of the electrostatic field, while LogAPBS allows to better identify the boundaries between regions of interactions favoring positive or negative charges (Figure 2:E,F).

## 3. Fields calculation and visualization

Field calculations can be launched from the command line and we provide examples for doing so. The input for the calculation comprises a structure file (PDB), the location of the center of the pocket, which, for example, can be given as a set of coordinates deduced from the presence of a ligand in a complex, and a radius for the pocket. It is important to notice that all calculations are then performed only on the receptor, ignoring the possible presence of a ligand already in the pocket. The algorithm defines a sphere centered at the given position and of the given radius *R* and builds a grid of preset mesh spacing inside. The mesh is chosen small enough to be able to follow smoothly changes in the fields (typically a fraction of an υ).

Field values are written in the form of 3D matrices of uniform sizes 2×R in each dimension, where each entry is the value of the given field at the spatial coordinate corresponding to the matrix indices. Values of the fields outside the sphere of interest are set to zero. In order to account for the physical volumes of the atoms of the receptors, field values for points in close proximity to pocket atoms are set to 0. The distance threshold for proximity is set to 2.5 υ for hydrogen bonding, for which atoms come in close contact because of the quasi-shared hydrogen, and to 3.0 υ for all other fields, for which atoms cannot come too close because of electronic shells repulsion. After this trimming, it can still happen that some points in space that are outside the actual pocket cavity are still included in the matrix. These are identified as clusters of points disconnected from all others. To remove these clusters, a graph traversal strategy [43] is employed, starting from the center of the sphere and visiting all untrimmed neighboring points; points that were not visited at the end of the traversal are further trimmed out. Each field grid is stored as a *.dx* or *.mrc* file, which can easily be imported with most molecular visualization programs, and then displayed through isosurfaces, represented in solid colors or as a mesh. A discussion on visualization strategies is presented in Supplementary Information, Section

## 2. Visualization plugins for the most common visualization software are available in the git repository

The typical time to compute a single grid is ∼ 1*s* for a pocket and ∼ 10 − 20*s* for a whole macromolecule, with a grid resolution of 0.25υ. The longest time is required to write the file in *.dx* format, which for a whole macromolecule with 10k atoms is ∼ 10 − 40*s*, while it is faster (≤ 10*s*) for the *.mrc* format. The calculation of the electrostatic field with APBS, which is done with the homonymous software, typically takes tens of seconds to minutes, depending on the system’s size: this is always the slower step in the pipeline to obtain the full set of fields.

To render the whole process of field calculations inside a known pocket straightforward and their visualization easily accessible, we implemented scripts to launch their calculations and to visualize them with UnityMol.

We also provide scripts for exploration in virtual reality and demonstrate its use in supplementary videos (see section Data availability). In the light of future applications in interactive docking for drug design, we tested field visualizations in virtual reality using an interactive docking mode under development. We used a stacking field visualization to guide precise positioning of a cyclic fragment in the scene, as illustrated in the supplementary video.

## 4. Benchmark Results

To assess the efficacy of the fields in locating favorable ligand positions, we designed a benchmark of 10 protein binding pockets and 10 RNA binding pockets with experimentally resolved structures in complex with a small ligand. We want to evaluate if the properties that emerge from the fields are in agreement with the known physical characteristics of the ligand. For example, when we compute a hydrophobicity field, a coherent result would be to find polar portions of the ligand to be inside of what we identify as the hydrophilic regions of the pocket, and non-polar portions to be in the hydrophobic parts. Similarly, for stacking we would expect to find aromatic portions of the ligand to be placed in regions of high stacking propensity of the stacking field, for hydrogen bonds we would expect to find donors/acceptors of the ligand inside the spheres of acceptors/donors of the pocket, and for electrostatics to find explicit charges of the ligand to occupy volumes with high electrostatic field values of opposite charge.

In this section we first report a few representative instances of pocket fields superposed to the experimental ligand, to visually illustrate the correspondence between fields’ high values (hot spots) and ligand position in the pocket. This analysis also illustrates the complexity of the fields observed in actual pockets, with features that could not have been guessed by simple observation of the pocket surface, as intrinsic to the pocket volume. Then, for each system, we build a pharmacophore-based model of the ligand using *pharmit* [44] to quantify the correspondence between the field properties and the ligand. We numerically check the overlaps between pharmacophores and the corresponding field counterparts.

We selected our benchmark pockets to cover a wide spectrum of interactions. Table 1 reports the systems chosen and the main modes of interaction between the pocket and the ligand identified from the literature and using the PLIP software [45]. For each system we show here only the field that matters the most for the interaction with the ligand, but all fields for all systems are shown in Supplementary Material, Section 4. A discussion on how to assess which field is most relevant is presented in Section 5.1.

**Table 1:** Protein benchmark (top), RNA benchmark (bottom) and most relevant interactions observed for each system as reported by PLIP. “ELE” means pure electrostatic interactions and gathers both salt bridges and metal complexation. “STK” gathers the *π*-stacking interactions. “HP” stands for hydrophobic (pseudo-)interactions. “HB” are hydrogen bonds.

| PDB ID | Molecule Type | Interaction |
| --- | --- | --- |
| 1BG0 | Arginine Kinase + ADP | ELE + HB |
| 1EBY | HIV-1 protease + inhibitor BEA369 | HB + HP |
| 1EHE | P450S + heme group (with Fe) | ELE + HB + HP |
| 1H7L | Glycosyltransferase SpsA + dTDP (and Mg) | ELE + HB + HP |
| 1IQJ | Human coagulation factor Xa + M55124 | STK + ELE + HB + HP |
| 1OFZ | Fungal lectin + beta-L-fucopyranose | ELE + HB + HP |
| 3DD0 | Carbonic anhydrase II + Ethoxzolamide | HP + HB + ELE |
| 3EE4 | Mn/Fe oxidase + myristic acid | HP + ELE |
| 5M9W | Thermolysin + inhibitor JC65 | HP + HB + STK + ELE |
| 6E9A | HIV-1 wt protease + PDB:J0S | HB + HP + ELE |
| 1AKX | RNA HIV + Argininamide | HB Acceptors/Donors |
| 1I9V | tRNA + Neomycin | HB Acceptors/Donors |
| 2ESJ | 16S RNA + Lividomycin A | HB Acceptors/Donors |
| 4F8U | bacterial ribosome RNAs + sisomicin | HB Acceptors/Donors |
| 5BJO | RNA aptamer + DFHO | HB Acceptors/Donors + STK |
| 5KX9 | FMN Riboswitch + Ligand 6YG | HB Donors + STK |
| 6TF3 | NAD<sup>+</sup> riboswitch + Cordycepin 5-triphosphate | HB Donors + STK |
| 7OAX | RNA aptamer + spermine | HB Acceptors/Donors |
| 7OAX | RNA aptamer + DMHBO | HB Acceptors/Donors + STK |
| 8EYV | Beetroot dimer RNA + DFHO | HB Donors + STK |

### 4.1. Selected examples

#### 4.1.1. Mn/Fe oxidase bound to myristic acid (PDB: 3EE4)

The Mn/Fe oxidase from *Mycobacterium tuberculosis* is an enzyme complex involved in the oxidative stress response and iron homeostasis within the bacterium [46]. This enzyme complex plays a critical role in detoxifying reactive oxygen species (ROS) and regulating intracellular iron levels to ensure bacterial survival and persistence within the host. PDB entry 3EE4 shows it in complex with myristic acid [47], a fatty acid that is often found as a component of phospholipids, triglycerides, and other lipid molecules.

The pocket for this system has been described as a narrow and hydrophobic cavity that widens at its internal end, leading to a larger volume where metal cations coordinate with the protein and with more H-bond possibilities [47]. The representation of the hydrophobic potential matches accurately with this description, as seen in Figure 3A. The aliphatic section of the ligand effectively overlaps with the mostly hydrophobic pocket, while the oxygen atoms at its extremity are placed in a hydrophilic region.

**Figure 3:**
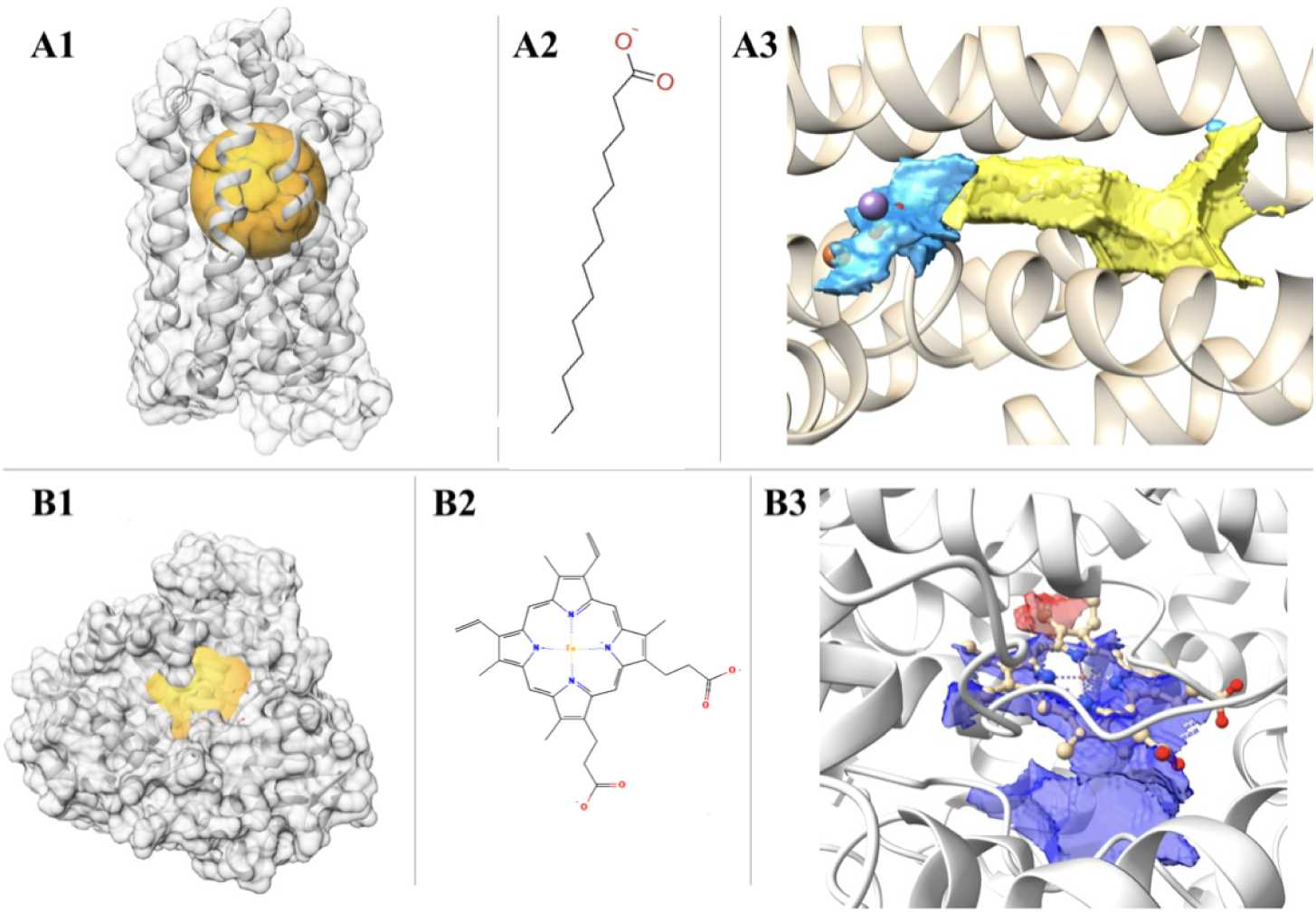
**A:** PDB system 3EE4: Mn/Fe oxidase bound to myristic acid. **B:** PDB system 1EHE: Monooxygenase cytochrome P450S bound to a heme group with iron. **A1) and B1)** representation of the whole protein structure and highlight (in orange) of the binding region where the fields are calculated; **A2) and B2)** 2D molecular graph of the ligand (A: myristic acid, B: heme group)); **A3) and B3)** the region where the field of interest is more intense (hydrophobic in yellow, hydrophilic in lightblue, positive/negative electrostatic in blue/red) superimposed to the ligands (balls and sticks/licorice representation). Protein structures shown with cartoon representation (B3: some protein atom surrounding the ligand also show in licorice).

#### 4.1.2. Monooxygenase cytochrome P450s bound to a heme group with iron (PDB: 1EHE)

Monooxygenase cytochrome P450s are a diverse superfamily of heme-containing enzymes found in all domains of life, including bacteria, archaea, fungi, plants, and animals [48]. They play essential roles in the oxidative metabolism of a wide range of endogenous and exogenous compounds, including drugs, fatty acids, steroids, and environmental pollutants. In PDB 1EHE, the oxygenase is bound to an heme group, which is notoriously capable of undergoing redox reactions, where the iron ion alternates between its ferrous (Fe2+) and ferric (Fe3+) oxidation states. This redox activity is essential for electron transfer in cytochrome proteins.

The binding pocket of this protein is characterized by a positively charged cluster of lysine and arginine residues, which plays an important role on how it interacts with the heme group ligand [49]. Indeed, the visualization of the electrostatic potential (Figure 3B3) reveals a positive electrostatic field in the binding pocket, which is ideal to accommodate four central electronegative nitrogens of the ligand, that then coordinate with the positive Fe ion. The positive field in the pocket, exhibits a hole in the location of the cation, therefore reinforcing its position.

Additional observations (shown in Supplementary Information and visible in the provided visualization files) include the stacking potential slightly overlapping with one of the aromatic groups, while the hydrogen bond potentials are displayed on top of the ligand’s heteroatoms. The hydrophobic main body of the ligand overlaps with a large region of hydrophobic potential, while the outside oxygen atoms are placed inside a hydrophilic section.

#### 4.1.3. Human coagulation factor Xa in complex with M55124 (PDB: 1IQJ)

Human coagulation factor Xa, often abbreviated as factor Xa, is a serine protease enzyme that plays a crucial role in the coagulation cascade, a series of biochemical reactions that lead to the formation of blood clots [50]. It is shown in complex with the inhibitor M55124 in the PDB entry 1IQJ.

The ligand’s aromatic pyridine group can be seen positioned in a significant stacking field region (Figure 4A3). Most of the pocket is hydrophilic (shown in SI), while the few hydrophobic sections neatly match the most hydrophobic parts of the ligand.

**Figure 4:**
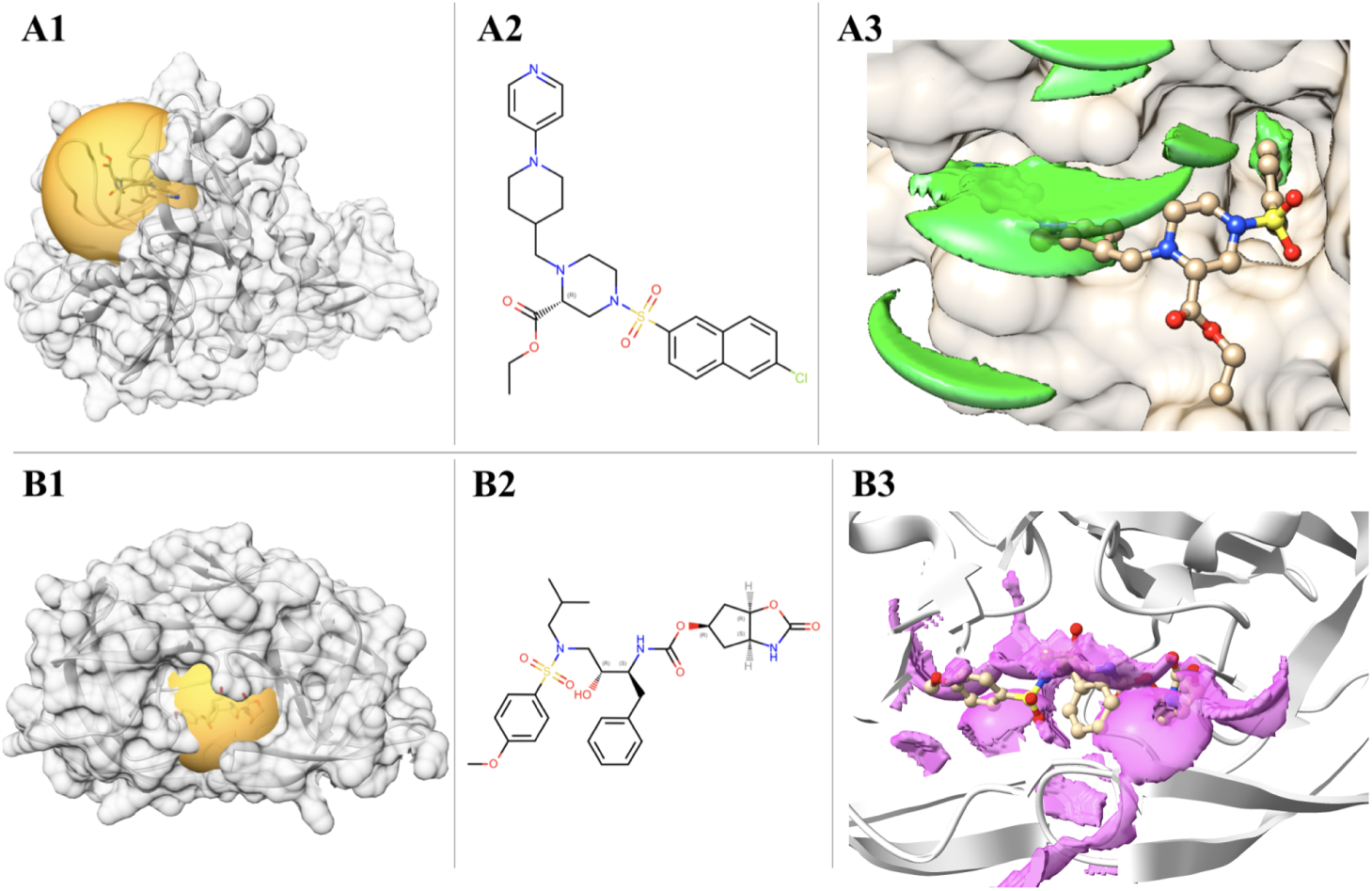
**A:** PDB system 1IQJ: Human coagulation factor Xa in complex with M55124. **B:** PDB system 6E9A: HIV-1 wild type protease bound to PDB:J0S. **A1) and B1)** representation of the whole protein structure and highlight (in orange) of the binding region where the fields are calculated; **A2) and B2)** 2D molecular graph of the ligand (A: M55124, B: PDB:J0S); **A3) and B3)** the region where the field of interest is more intense (stacking in green, HBA in magenta) superposed to the ligands (balls and sticks representation).

#### 4.1.4. HIV-1 wild type protease bound to PDB:J0S (PDB: 6E9A)

The HIV-1 protease plays a critical role in the viral life cycle by cleaving the viral polyprotein precursor into functional, mature proteins necessary for viral replication and assembly [51]. In [51], the authors report a plethora of new potent HIV-1 protease inhibitors and in PDB entry 6E9A the protease is shown in complex with the chemical compound corresponding to the PDB entry J0S.

The pocket for this protein is reported to have a high binding affinity with its ligands, which can be attributed to the possibility of creating multiple hydrogen bonds and hydrophobic interactions [51]. The hydrogen bond acceptor field visualized in Figure 4B3 overlaps nicely with the expected oxygens and nitrogens of the ligand (this can be visualized more easily with the provided visualization files). The hydrophobic and hydrophilic regions (see SI) appear to coincide to their counterparts in the ligand.

#### 4.1.5. FMN Riboswitch in complex with ribocil (PDB:5KX9)

Riboswitches are RNA molecules involved in regulation of gene expression, mostly found in bacteria. Their mode of action involves a large conformational rearrangement, where the full structure is modified at the level of base pairing, and therefore for the full three-dimensional structure as well. This rearrangement is triggered by a ligand and an inhibition mechanism consists of designing antagonists that bind in place of the natural ligand, thereby preventing its interaction with the target complex such that the riboswitch conformational transition is halted. The flavin mononucleotide riboswitch (FMN) is found in Escherichia coli and the molecule ribocil has been found experimentally to inhibit its function [52].

According to the literature, the interaction in this complex is driven by stacking and hydrogen bonding. From our analysis we can clearly observe in Figure 5 how the stacking field superposes almost perfectly with the position of the rings of the ligand inside the pocket. The hydrogen bond acceptor field also exhibit a relevant superposition with the locations of the Oxygens and Nitrogens of the ligand.

**Figure 5:**
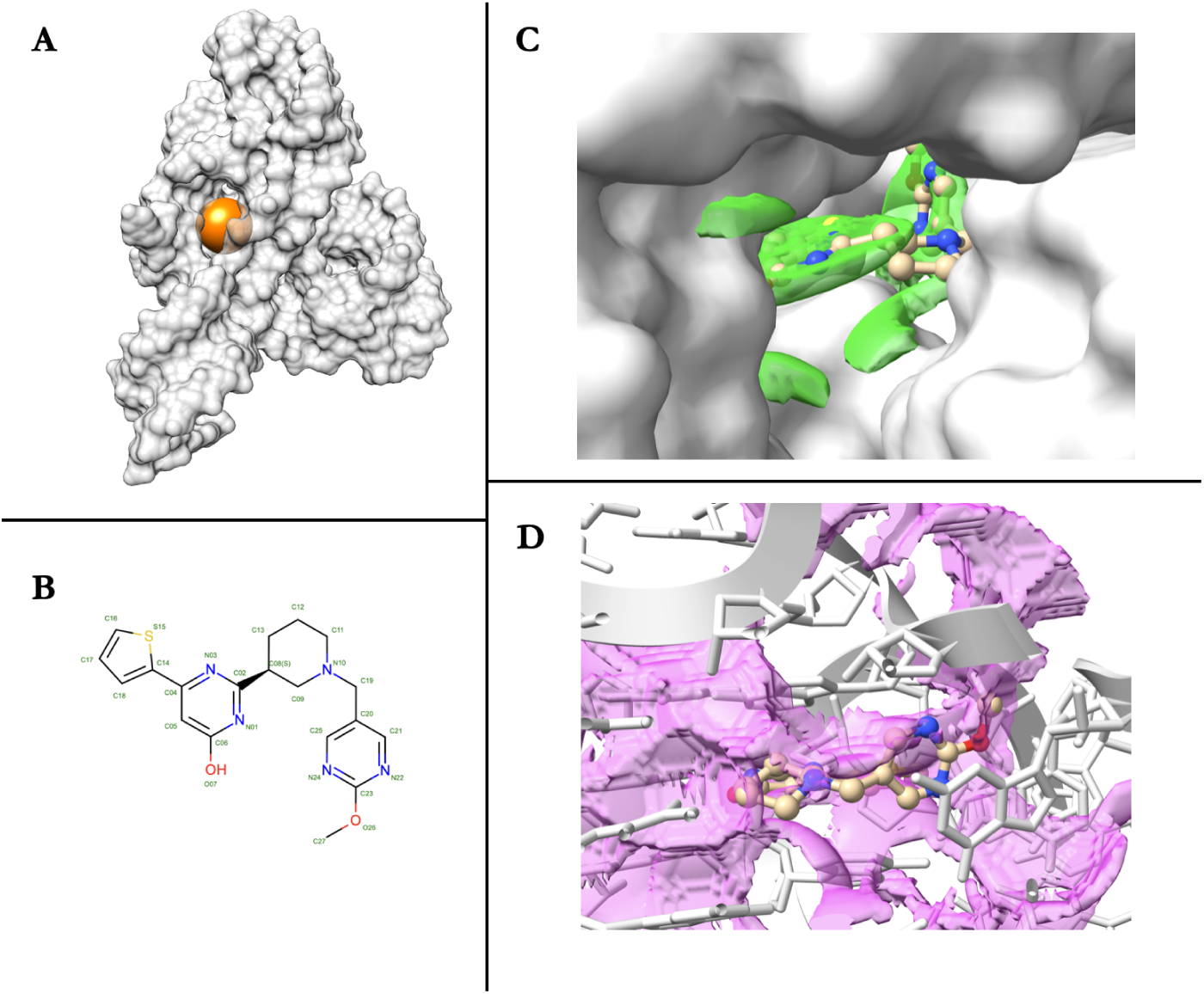
**A:** PDB system 5KX9: FMN Riboswitch in complex with ribocil. **B:** Chemical structure of the ligand ribocil (PDB:6yG). **C:** Representation of the stacking field (green) inside the pocket superposed to the ligand. **D:** Representation of the HBA field (magenta) inside the pocket superposed to the ligand.

#### 4.1.6. Homo sapiens ribosomal decoding site

As a last example, we report a case where multiple ligands are known and experimentally solved for the same molecule, even in the same binding site. The target is the eukaryotic ribosomal decoding site, and the PDB entries we used are 2G5K (HOLO form, bound to apramycin, or AM2, in two different binding sites) and 2O3V (HOLO form, bound to a synthetic compound named NB33). We report the structures and fields of 2G5K and 2O3V together with the ligands AM2 and NB33 (Fig.6A).

**Figure 6:**
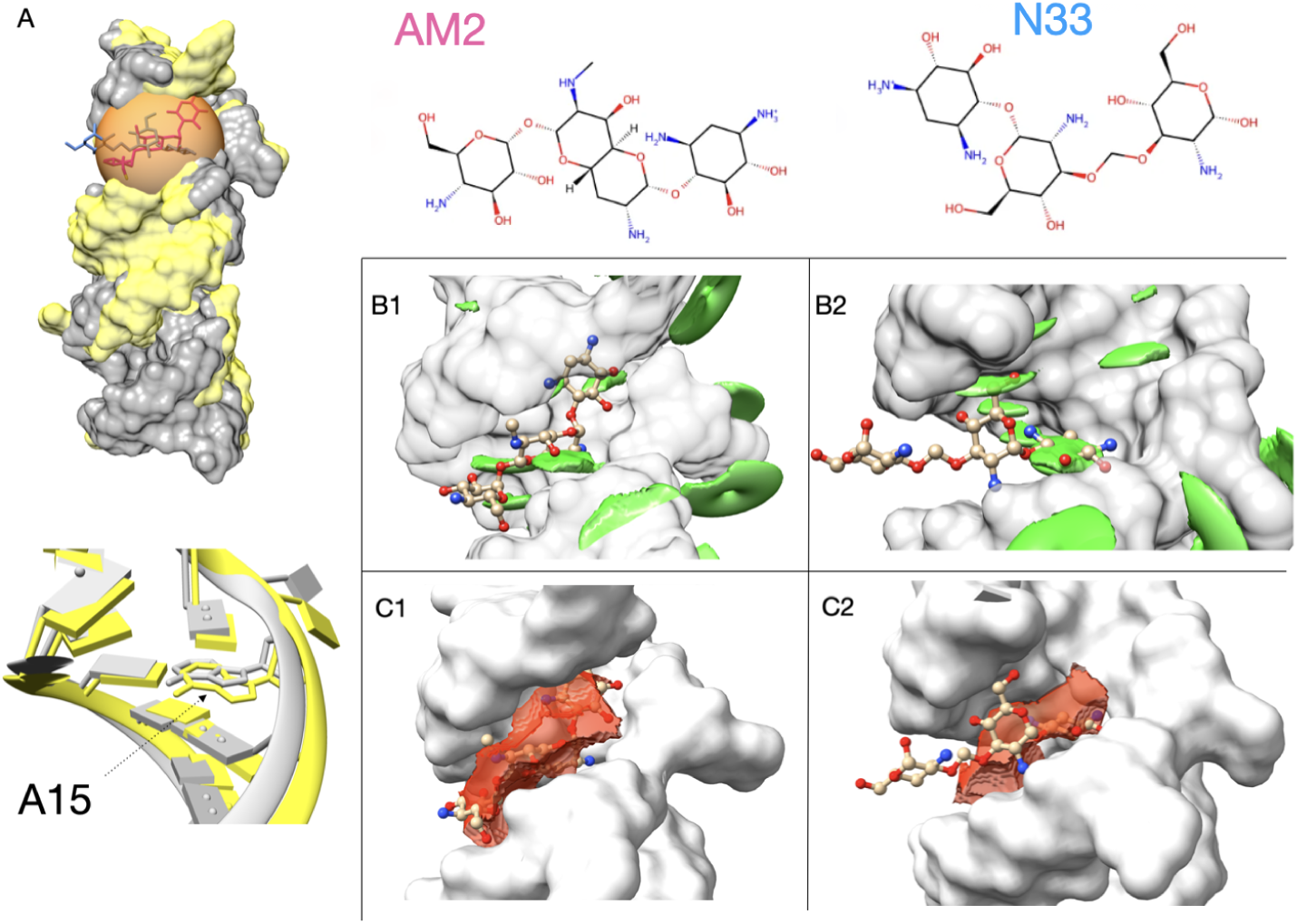
A: 3D structure of the *Homo sapiens* ribosomial decoding site in complex with 2 different ligands: NB33 (blue/gray, from PDB 2O3V), AM2 (magenta/yellow, from PDB 2G5K). Details of the position adopted in the two structures by base A15 are shown in the bottom of the panel. Stacking (B1 and B2) and electrostatic (C1 and C2) fields of the binding pockets superposed with the ligands. Protonation state of the ligands at physiological pH has been obtained following [53].

The secondary structure of the RNA system is the same in both the structures, but interestingly one can notice that the unpaired bases have different dispositions in the 3D space. One of these differences is crucial for the interaction with the ligands: in RNA-NB33 the adenine in position 15 (with the atoms and bonds of the base explicitly represented in Fig. 6A, lower image) is slightly more exposed towards the binding site than in RNAAM2, allowing the formation of a stable stacking interaction with one ring of NB33 (Fig. 6B2) which is absent in the RNA-AM2 complex (Fig. 6B1). In contrast, AM2 appears to use more the electrostatic interaction, as it fully sits in the region where the electrostatic field is highest, which is coherent with its protonation state. Its position is possibly also further stabilized by the presence of a magnesium ion at the edge of the pocket. However, this interaction appears to be less specific than that of NB33, as the electrostatic high field is diffused in the whole pocket. This observation is in line with the discussion made in [54], where the authors not only report the crystal structures of these complexes, but also demonstrate an increased binding affinity of NB33 to the eukaryotic A-site with respect to the prokaryotic counterpart, and a selectivity of NB33 to bind only one of the 2 possible pockets found in the RNA system, differently from AM2.

### 4.2. Overlaps of fields with known pharmacophores

To quantify in a general manner what is observed through visual inspection, for the ligands present in the benchmark structures, we compared the spatial regions where the fields assume high (absolute) values with the pharmacophore models generated by *pharmit* [44]. Regions with lowor zerointensity fields were filtered out by retaining only those points with values that deviate by a maximum of 2 sigma from the peak value of the individual Gaussian function. For the electrostatic fields, a threshold of ±20*k_B_T/e_c_* was chosen by visual inspection to highlight the main features of the field. The pharmacophores (PPs) of each type modeled by pharmit were compared with the filtered fields of their counterpart, remembering that the fields refer to the receptor, while the pharmacophores refer to the ligand. Therefore we have to compare stacking site PP with stacking field, hydrophobic field with hydrophobic PP, but negative charge site PP with positive APBS field, hydrogen bond donor field with hydrogen bond acceptor PP, etc. To make this comparison, for each system we first evaluate the interactions between the ligand and the receptor using the PLIP software [45] to identify the PPs of the ligands that are actually involved in the interaction, and then check if there is an overlap between the PPs and the corresponding field. We count the interaction if there is a spatial overlap between the filtered fields and a sphere around the PP coordinates whose radius is chosen in agreement with the literature [55]: R*_HBA_* = 0.5 υ, R*_HBD_* = 1.0 υ, R*_ST K_* = 1.0 υ, R*_HP_* = 1.0 υ, R*_ELE_* = 1.0 υ. No comparison has been made with the *hydrophilic* field, since hydrophilic pharmacophores don’t exist.

Table 2 summarizes the number of PPs that overlap with the filtered fields compared to the interactions predicted by PLIP. The comparison between fields, pharmacophores and PLIP predictions, however, requires some care. In a few cases, the number of pharmacophores overlapping with the fields is higher than the number of interactions identified by PLIP, suggesting the existence of interactions not captured by PLIP criteria, possibly due to strict cutoffs used by PLIP to identify hydrogen bonds and stacking, which could result in PLIP missing a few meaningful interactions. An example of interaction not accounted for by PLIP but for which we have an agreement between pharmacophores and field is presented in Figure 7. In other cases pharmit does not predict pharmacophores where PLIP detects an interaction. This occurs for example for 6E9A and 3EE4 HP where the fields agree with PLIP but the pharmacophores are missing. We note that PLIP detects only one hydrophobic interaction for RNA systems. The combination of pharmit and the fields instead detects a substantially higher number of these interactions. The discrepancy is due to the fact that PLIP attributes only one property to the interaction, either stacking or hydrophobic, while pharmit accounts for both. Since most hydrophobic interactions in RNA arise from the aromatic rings, PLIP attributes them mainly to stacking.

**Figure 7:**
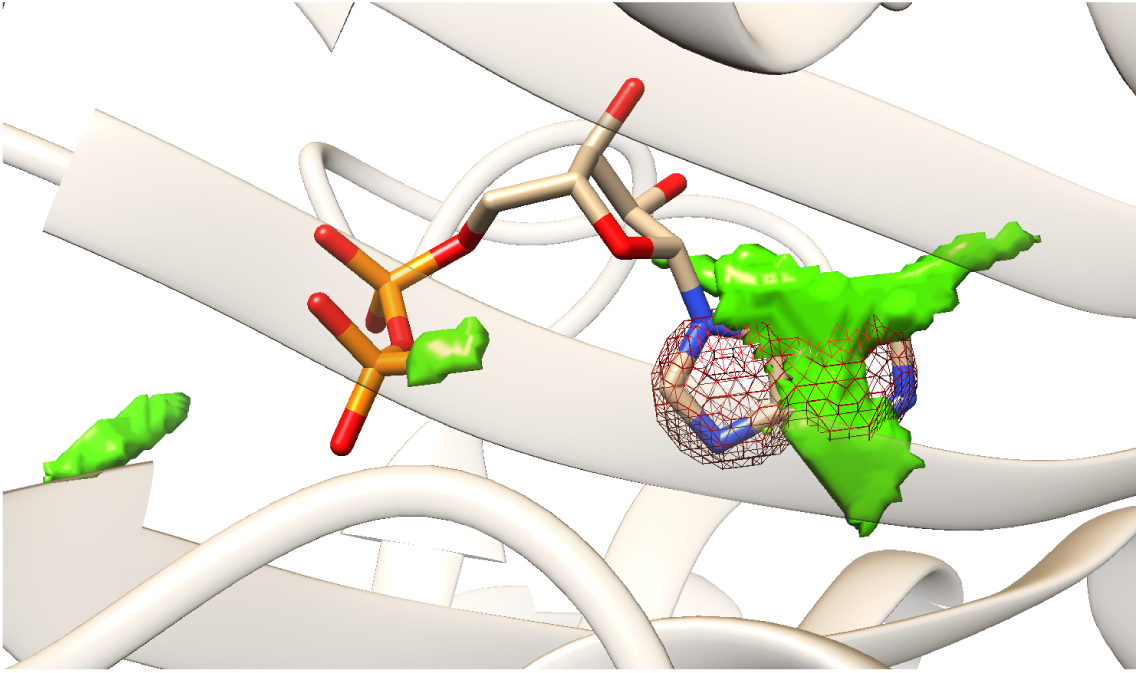
Zoom around the ligand ADP in PDB 1BG0, showing the filtered stacking field in green (as described in the text) and the relative pharmacophores’ field in brown, as a mesh grid. In this case, 2 stacking PP sites are overlapping with the field although one to a lesser extent than the other. In this case, PLIP detects 1 stacking interaction between the protein and ADP.

**Table 2:** Table representing the number of interactions detected by the fields, given by counting the pharmacophores of a given species that overlap with the corresponding interaction field. Each element in the table is in the format *# of overlaps* (*# of total pharmacophores*) *# of PLIP interactions*. Rows labeled “TOT” indicate the number of PP overlapping with fields that match those predicted by PLIP.

|  | HBA |  | HBD |  | HP |  | NEG |  | POS |  | STK |  |
| --- | --- | --- | --- | --- | --- | --- | --- | --- | --- | --- | --- | --- |
| **1bg0** | 8(13) | 3 | 3(3) | 2 | 1(2) | 0 | 2(2) | 2 | 0(0) | 0 | 2(2) | 1 |
| **1eby** | 2(8) | 2 | 6(6) | 3 | 4(4) | 4 | 0(0) | 0 | 0(0) | 0 | 0(4) | 0 |
| **1ehe** | 4(6) | 1 | 0(0) | 0 | 8(8) | 8 | 2(2) | 2 | 0(0) | 0 | 0(2) | 0 |
| **1h7l** | 2(11) | 3 | 1(2) | 1 | 2(2) | 2 | 0(2) | 1 | 0(0) | 0 | 1(1) | 0 |
| **1iqj** | 1(5) | 0 | 0(0) | 0 | 4(5) | 2 | 0(0) | 0 | 0(0) | 0 | 1(3) | 1 |
| **1ofz** | 2(5) | 2 | 3(4) | 1 | 1(1) | 1 | 0(0) | 0 | 0(0) | 0 | 0(0) | 0 |
| **3dd0** | 2(4) | 1 | 1(1) | 0 | 3(3) | 1 | 0(0) | 0 | 0(0) | 0 | 0(2) | 0 |
| **3ee4** | 0(2) | 0 | 0(0) | 0 | 5(5) | 7 | 0(1) | 0 | 0(0) | 0 | 0(0) | 0 |
| **5m9w** | 3(11) | 3 | 6(8) | 2 | 2(2) | 2 | 0(2) | 1 | 0(0) | 0 | 0(1) | 0 |
| **6e9a** | 2(6) | 2 | 3(3) | 1 | 4(5) | 7 | 0(0) | 0 | 0(0) | 0 | 0(2) | 0 |
| **TOT** | 16/17 |  | 10/10 |  | 29/34 |  | 4/6 |  | 0/0 |  | 2/2 |  |
| **1akx** | 1(1) | 1 | 2(4) | 1 | 1(1) | 1 | 0(0) | 0 | 1(1) | 1 | 0(0) | 0 |
| **1i9v** | 0(19) | 0 | 5(13) | 2 | 1(1) | 0 | 0(0) | 0 | 0(0) | 0 | 0(0) | 0 |
| **2esj** | 2(23) | 4 | 9(15) | 3 | 1(1) | 0 | 0(0) | 0 | 0(0) | 0 | 0(0) | 0 |
| **4f8u** | 2(12) | 3 | 8(8) | 3 | 3(3) | 0 | 0(0) | 0 | 0(0) | 0 | 0(0) | 0 |
| **5bjo** | 2(4) | 1 | 2(2) | 1 | 4(4) | 0 | 0(0) | 0 | 1(1) | 1 | 1(1) | 1 |
| **5kx9** | 1(6) | 1 | 0(0) | 0 | 3(3) | 0 | 0(0) | 0 | 1(1) | 0 | 1(1) | 1 |
| **6tf3** | 3(12) | 2 | 2(2) | 0 | 2(2) | 0 | 0(2) | 0 | 0(0) | 0 | 2(2) | 2 |
| **7oax0** | 0(4) | 0 | 4(4) | 1 | 3(3) | 0 | 0(0) | 0 | 0(0) | 0 | 0(0) | 0 |
| **7oax1** | 2(7) | 0 | 2(2) | 0 | 5(5) | 0 | 0(0) | 0 | 0(0) | 0 | 2(2) | 2 |
| **8eyv** | 1(4) | 0 | 2(2) | 0 | 4(4) | 0 | 0(0) | 0 | 1(1) | 0 | 1(1) | 1 |
| **TOT** | 9/12 |  | 11/11 |  | 1/1 |  | 0/0 |  | 2/2 |  | 7/7 |  |

Results presented in the table are therefore to be interpreted with care and are mainly qualitative (also because of the approximations made in the filtering parameters for the fields, the radii for the pharmacophores, and, to some extent, for the cutoff of the initial trimming). However, it is clear that we obtain an overall excellent correspondence between the regions of high field values and the pharmacophores involved in the ligand’s interaction with the receptor.

## 5. Binding pocket characterization

Since computing fields with SMIFER is very fast, we can easily compute the fields of a system as a whole, not limiting to a predefined binding pocket. This opens the road to a fuller characterization of the properties of a pocket. In general, absolute values of the fields inside the pocket are not very meaningful since they vary significantly from system to system. However, values relative to the fields around the rest of the system outside the pocket are more informative as they help highlighting the regions where the fields are strongest, which could favor the interaction with a binding partner. The following two applications are built on the comparison of a field inside a pocket relative to the field all around the system.

### 5.1. Qualitative assessment of field relevance

Even though we typically compute all fields at once, they don’t all bear the same relevance for a specific system and for a binding pocket. In order to get an insight on the importance of a field *i* for a given pocket *p*, we can compute a relevance score (*s^p^*) as follows. Having identified the volume *V_p_* of a pocket (either by proximity with a known ligand or by predictions), we compute the ratio of the integral of the values of the field in the pocket and of the integral of the field over the whole volume defined by the grid for the whole system. The relevance score is therefore computed as:

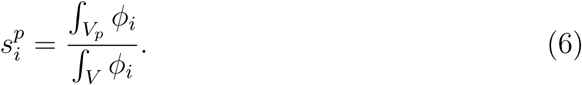

where *ϕ_i_* is the field of interaction *i*, and *V* is the overall volume used to compute the field for the whole system. The idea behind this simple score is that a field is more relevant if it is concentrated in the pocket as a ligand fulfilling this property might be most likely to interact favorably at that location.

If we want to compare different systems, for which the overall integral of the fields varies, we can introduce a normalized relevance score, named relative relevance score as:

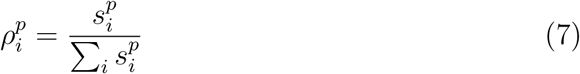

For each system in the benchmark the contribution of each field is shown in a bar plot (Figure 8). For proteins the fields considered are : electrostatics, HBA, HBD, stacking, hydrophobic and hydrophilic, while for RNA we exclude hydrophobic and hydrophilic in this calculation as the hydrophobic field is redundant with stacking and hydrophilic is almost always absent. Results on the benchmark systems qualitatively agree with the observed ligand properties in the pocket. For RNA some stacking and some HB are always present, but those systems for which stacking is highest are indeed found in complex with ligands that use stacking as a mode of interaction, and the same holds true for HB interactions. For proteins, for some systems it is clear that one field dominates, especially when electrostatics or hydrophobicity are high, while for others the contribution seems to be more evenly distributed between the various interactions, which is in agreement with the different interactions detected by PLIP for these systems (see Table 1).

**Figure 8:**
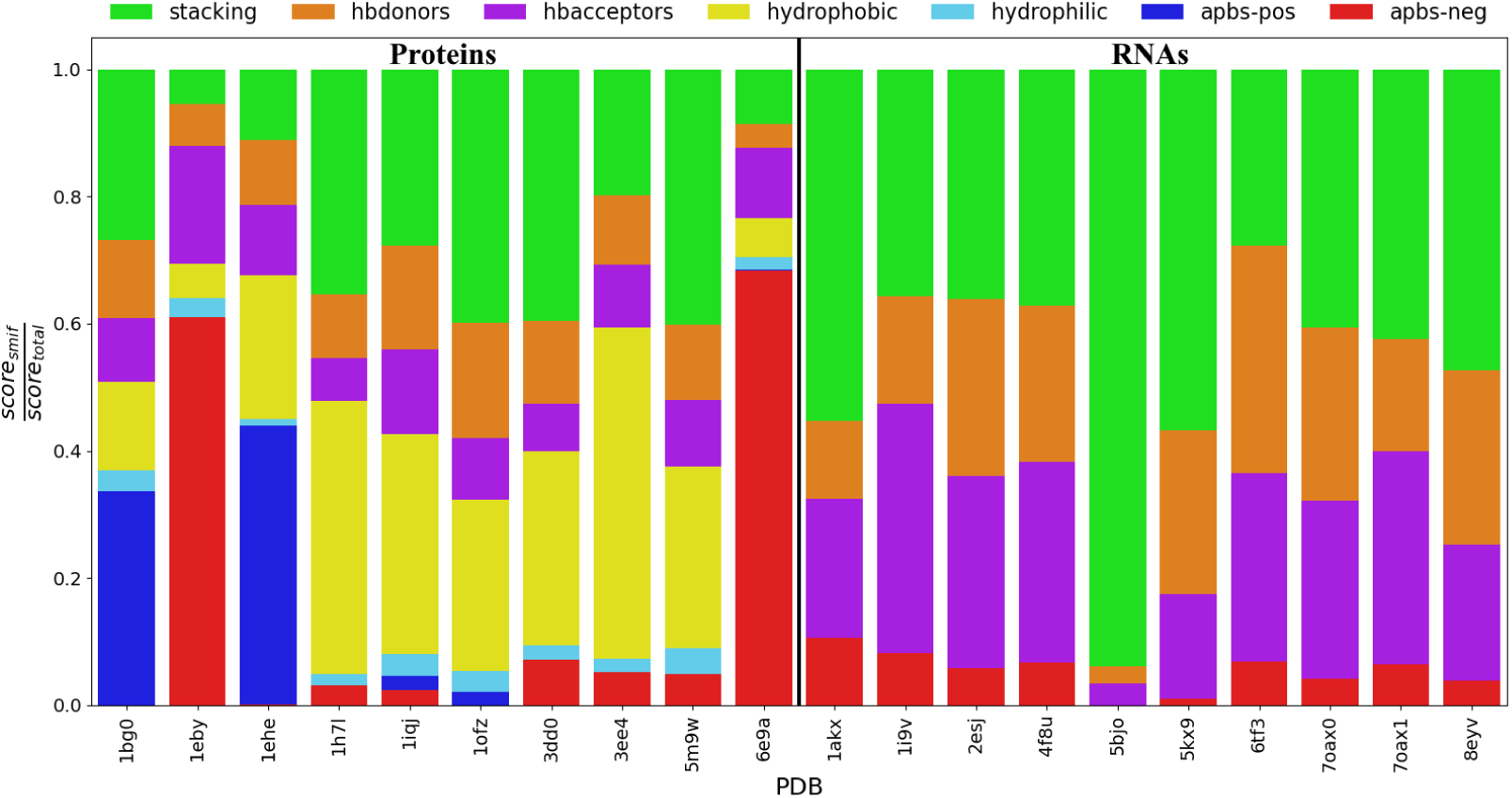
For each benchmark system the barplot shows the relative relevance score *ρ^p^*. The first 10 systems are proteins for which hydrophobic and hydrophilic fields are included in the calculation, while they are excluded for the second 10 systems which are RNAs.

#### RNA binding pockets

The relevance scores of the fields inside a given pocket can be used to compare the nature of different pockets within a given system. For this purpose, we can define a pocket score as the sum of the relevance scores of each field inside a pocket normalized by the volume of the pocket *S_p_* = *s^p^/V_p_*, effectively giving a density of fields inside the pocket.

This approach can be particularly interesting for RNA molecules, for which identifying and comparing possible pockets is still an open challenge. We show here an example of how fields can be used to classify and to gather an understanding of the properties of possible RNA pockets using the riboswitch example discussed earlier (PDB ID 5KX9). This system is found in complex with a ligand that sits in the center junction, in a deeply buried pocket. However, in some systems reported in RNA/ligand databases (such as HARIBOSS), ligands are found also in shallow pockets closer to the surface. It would therefore be possible that the riboswitch could eventually have ligands binding elsewhere, possibly making use of different kind of interactions with respect to the experimental ligand.

We use the programs fpocket [56] and fpocketR [57] to identify regions of the riboswitch that could be pockets based on geometric properties, and we then score these putative pockets based on the relevance of the fields inside. Excluding absurdly large pockets, the two programs propose 10 pockets, but pockets 1 and 2 overlap quite significantly and can be considered the same pocket in practice. This is also the location of the ligand in the experimental complex. Figure 9 shows the 10 pockets found for this system and the pocket score *S_p_* for each of them, keeping track of the contribution of each field. Clearly, for this system the pocket buried inside the structure (pockets 1 and 2) is the most favorable one, with a very high stacking potential, followed by high HB potentials. However, other possible pockets could have interesting properties and eventually bind to other ligands. For a small ligand with good stacking properties pocket 4 and 5 could be an option, while extended ligands with high HB capabilities and no stacking could accommodate in the groves in pockets 3, 7 and 8, as it is found in some other systems, even though these molecules might not have drug-like properties.

**Figure 9:**
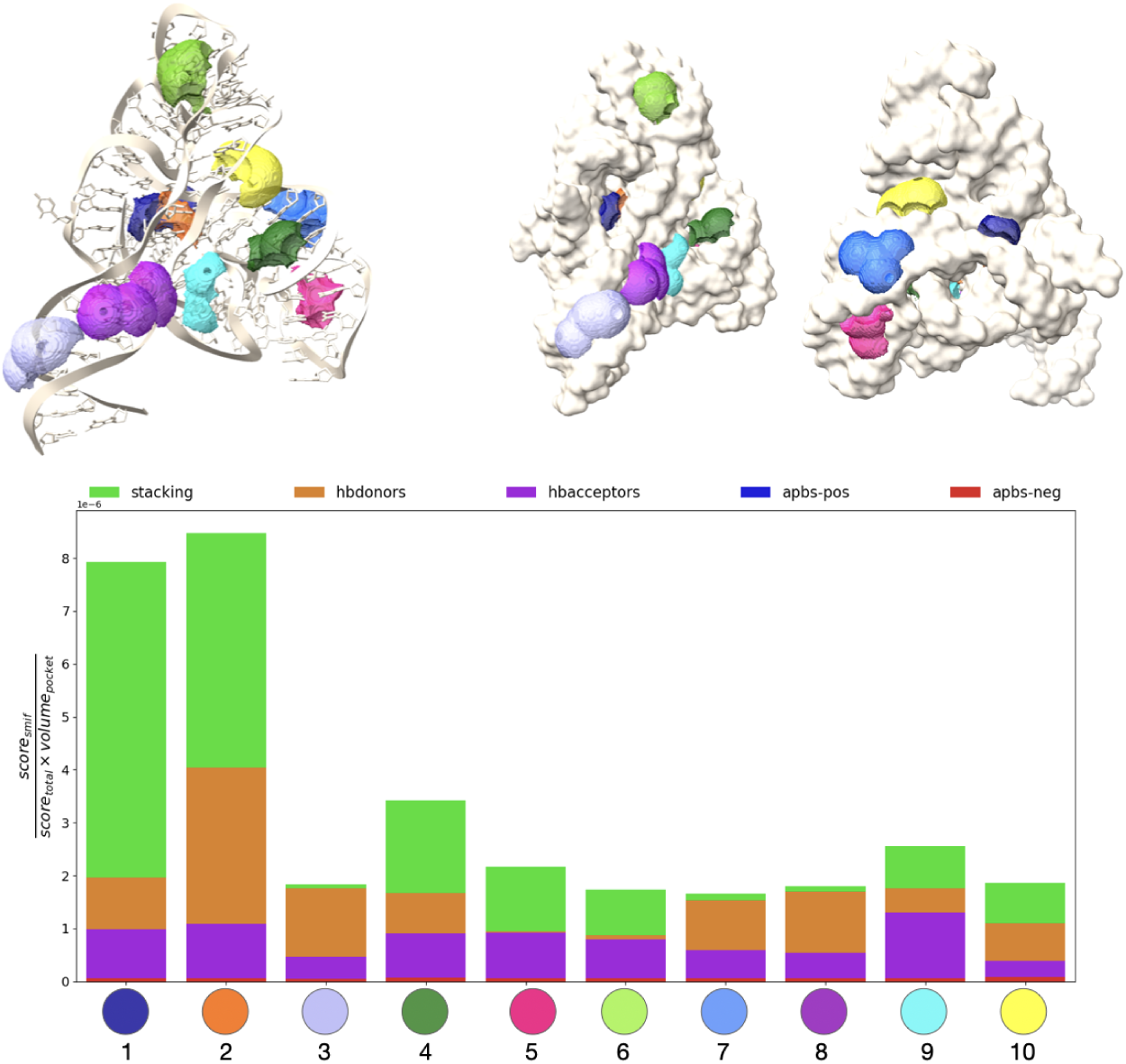
Top: pockets identified by fpocket and fpocketR for the riboswitch 5KX9. On the right pockets are visualized together with the surface of the molecule from two different view points for a better insight on their buriedness. Bottom: The pocket-scores of the various fields in a given pocket are presented for each pocket. The higher the score, the stronger the binding that can putatively be achieved.

This analysis, even though still at its preliminary stage, is a proof of principle that fields can provide insightful information to score pockets for their overall likelihood based on the overall strengths of the fields, but also classify them for the different kinds of interactions they can accommodate, in a very interpretable manner. It is interesting to notice that the ranking of our scores agrees with the results of fpocketR (pockets 1 and 2), which is a very recent pocket predictor trained specifically for RNA molecules and drug-like ligands. Our analysis adds to it the insight of the contributions of different kinds of interactions.

## 6. Perspectives on large systems

Having assessed the usefulness of computing fields to understand the physical properties of binding pockets, we tested field calculations on whole systems to evaluate the extent to which fields might be useful to characterize the properties of macromolecules more in general.

### Membrane proteins hydrophobicity

The Human GABA_A_ receptor alpha1-beta2-gamma2 (PDB ID 6X3V, protein only) is a transmembrane protein containing an ion channel. We investigate in particular the hydrophobic and hydrophilic fields as we expect to detect differences in these fields between the regions that are embedded in the membrane and the parts exposed to the solvent, as well as between the inside of the ion channel and the outside. Our hydrophobic/hydrophilic field is shown in Figure 10 for the whole systems (A) and for slices between planes parallels to the membrane (B). We can observe how the outside of the molecule offers a hydrophobic environment around the transmembrane helices while the head of the molecule is mainly hydrophilic. In particular the entrance of the ion channel is hydrophilic. As we move down the molecule following the channel we can observe that the environment inside the channel remains always hydrophilic (B1-B4), while the external part becomes hydrophobic in correspondence of the transmembrane helices (B2 and B3). The use of fields offers a very clear view of the hydrophobic and hydrophilic regions around the molecule which is hardly achievable by typical visualization coloring the surface using a hydrophobic/hydrophilic scale.

**Figure 10:**
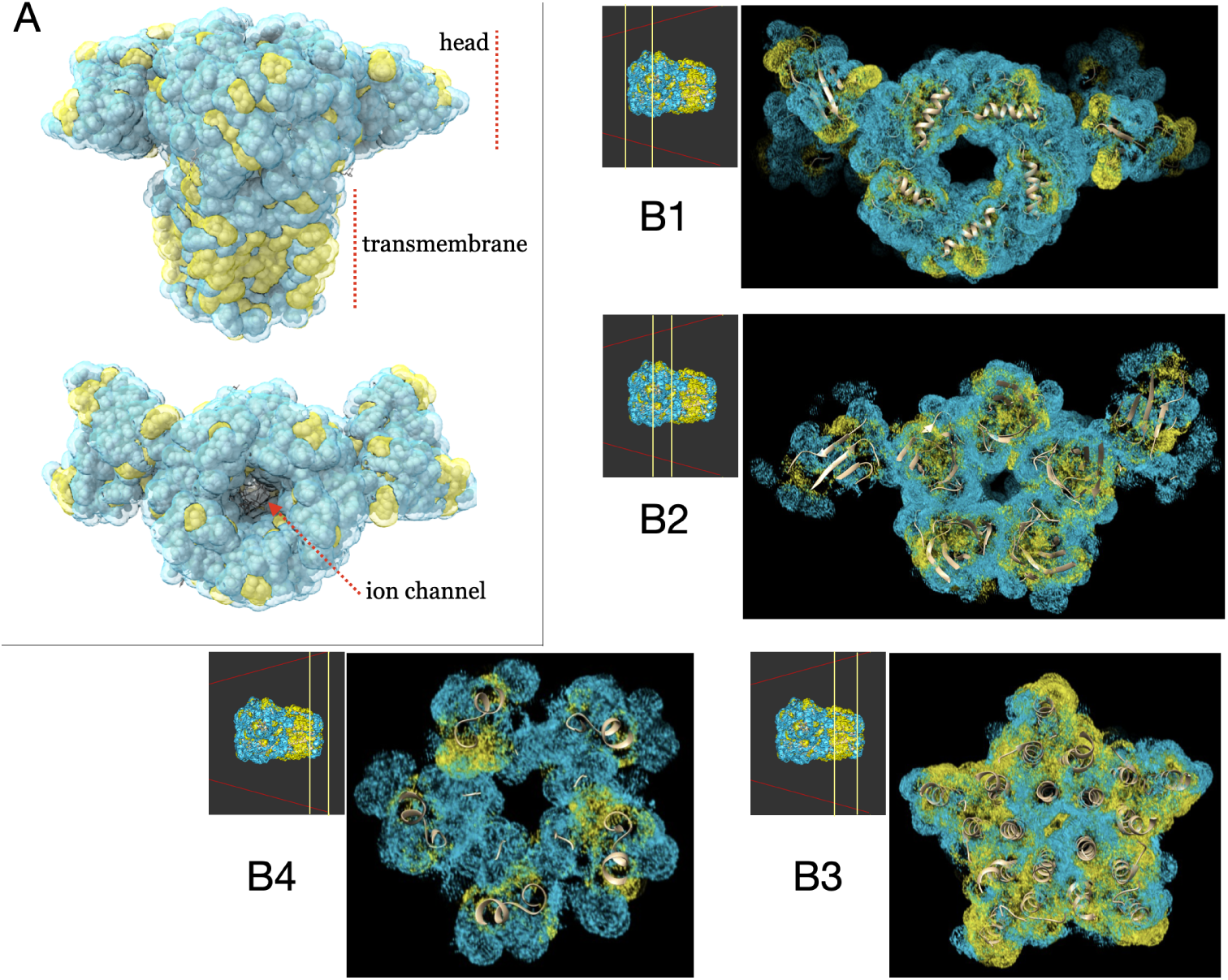
Visualization of the hydrophobic (yellow) and hydrophilic (light blue) fields for the GABA_A_ receptor. A: Visualization of the fields for the whole system (side and top view), B: visualization of the fields for slices as we move across the channel from the head to the transmembrane region as shown in the smaller quadrant (the region shown is in between the two yellow lines).

### Large systems interfaces

As last application, we illustrate the analysis of the interface of a large complex consisting of the Internal Ribosomal Entry Site (IRES) of Hepatitis C bound to the 40S ribosomal subunit. Cryo-EM experiments show two distinct complexes one in which the IRES is bound but closed on itself (PDB ID 7SYG), and a second one in which the IRES stretches across the ribosomal subunit (PDB ID 7SYP) in an open form [58]. We want to use our fields to characterize the binding interfaces and gather insights on the binding modes of the two IRES configurations.

To achieve this goal we want to identify what fields are ”used” in the binding, *i. e.* find those that actually form an interaction with the corresponding partner in the other molecule. For example, when considering a stacking field, we want to see what regions of high field values match the position of aromatic rings of the other molecule. Similarly, for hydrogen bonds, for the donor field we want to see what positions are matched with an acceptor atom of the binding partner. For the electrostatics, we will compare the field with a charge density field of the binding partner. In practice, to compute what we will call the ”interface fields”, we define a new class of fields that are just the occupation of the binding partners and we compute the voxel-wise product between the statistical fields of one molecule and the occupation field of the partner. Occupation fields are simply spheres centered around the binding partner elements (HB donor atoms, HB acceptor atoms, aromatic rings, hydrophobic and hydrophilic atoms), with radius 1υ. The only outlier is the charge density, for which we consider normalized gaussians centered around the sites of partial charges with width 1υ and weighted by the value of the charge. These choices were mainly dictated by the visual rendering, since at these level we are not interested in a quantitative characterization of the interactions. The mathematical definition of the occupation fields follows:

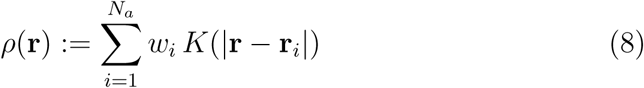

where *K*(**r**) is a generic kernel that can be modeled either as a Heaviside step function defined over a sphere or a normalized Gaussian function with *µ* = 0υ and *σ* = 1υ; *w_i_* is 1 for the sphere-like occupation field while it corresponds to the partial charge of the atom *i* in the charge density case.

We perform this analysis on the open and closed conformations of IRES: results are illustrated in Fig.11 and Fig.12 and in the video provided as supplementary material. Figure 11 provides a detailed analysis of the electrostatic interactions, where both open and closed conformations exhibit regions of attractive (green) and repulsive (orange) electrostatic patches. The iso-value chosen for the figure corresponds to an energy density of ∼ ±0.2 kT/υ^3^.

**Figure 11:**
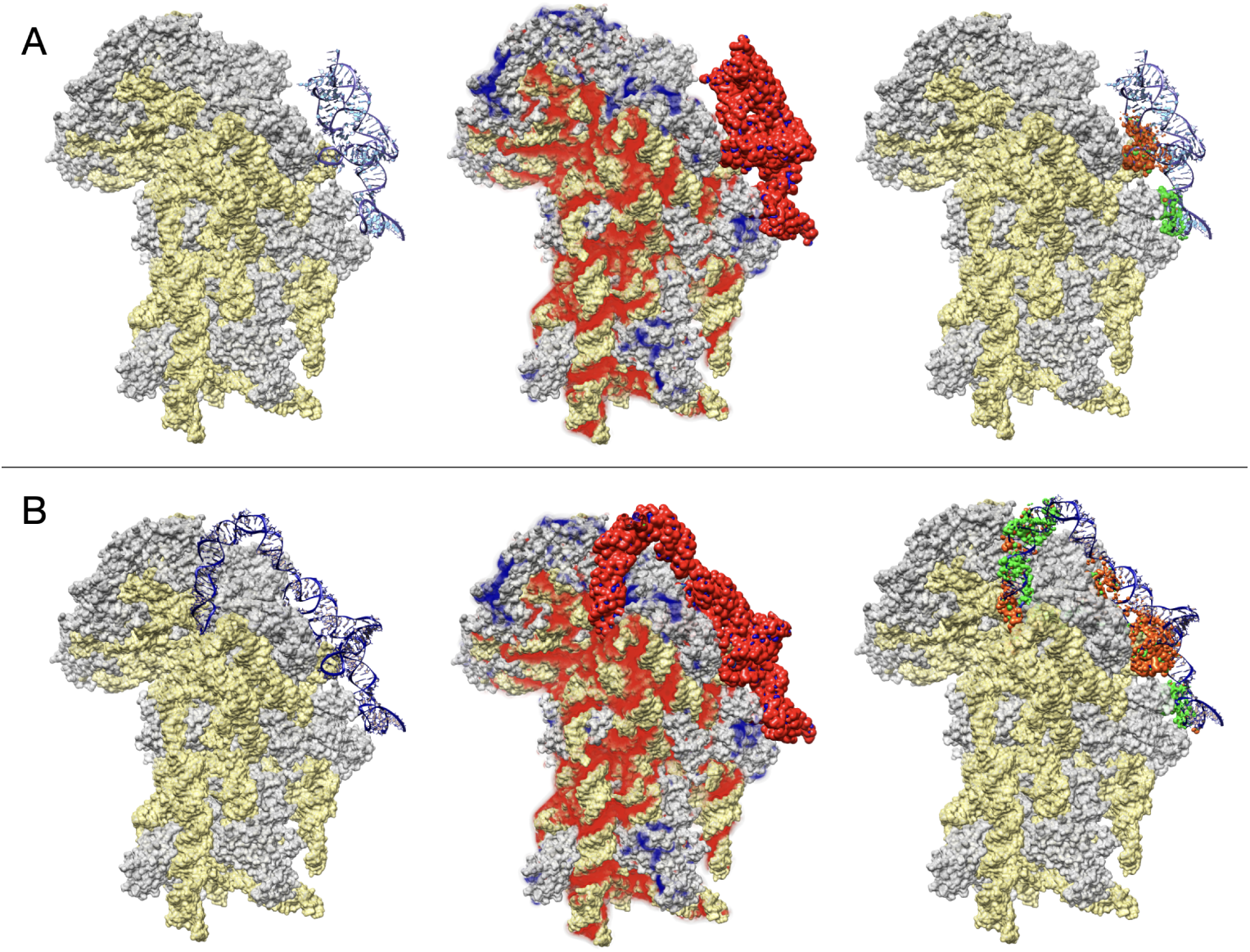
A: Closed IRES configuration, B: open IRES configuration. The ribosomal subunit is shown in gray (protein) and yellow (RNA). Left: structures from the CryoEM with the IRES shown in blue. Center: electrostatic field computed for the ribosomal subunit and the charge density computed for the IRES (red: negative field/charges, blue: positive field/charges). Right: results of the product of the electrostatic field by the charge density. We report in green regions of favorable/attractive interactions and in orange the regions of unfavorable/repulsive interactions.

**Figure 12:**
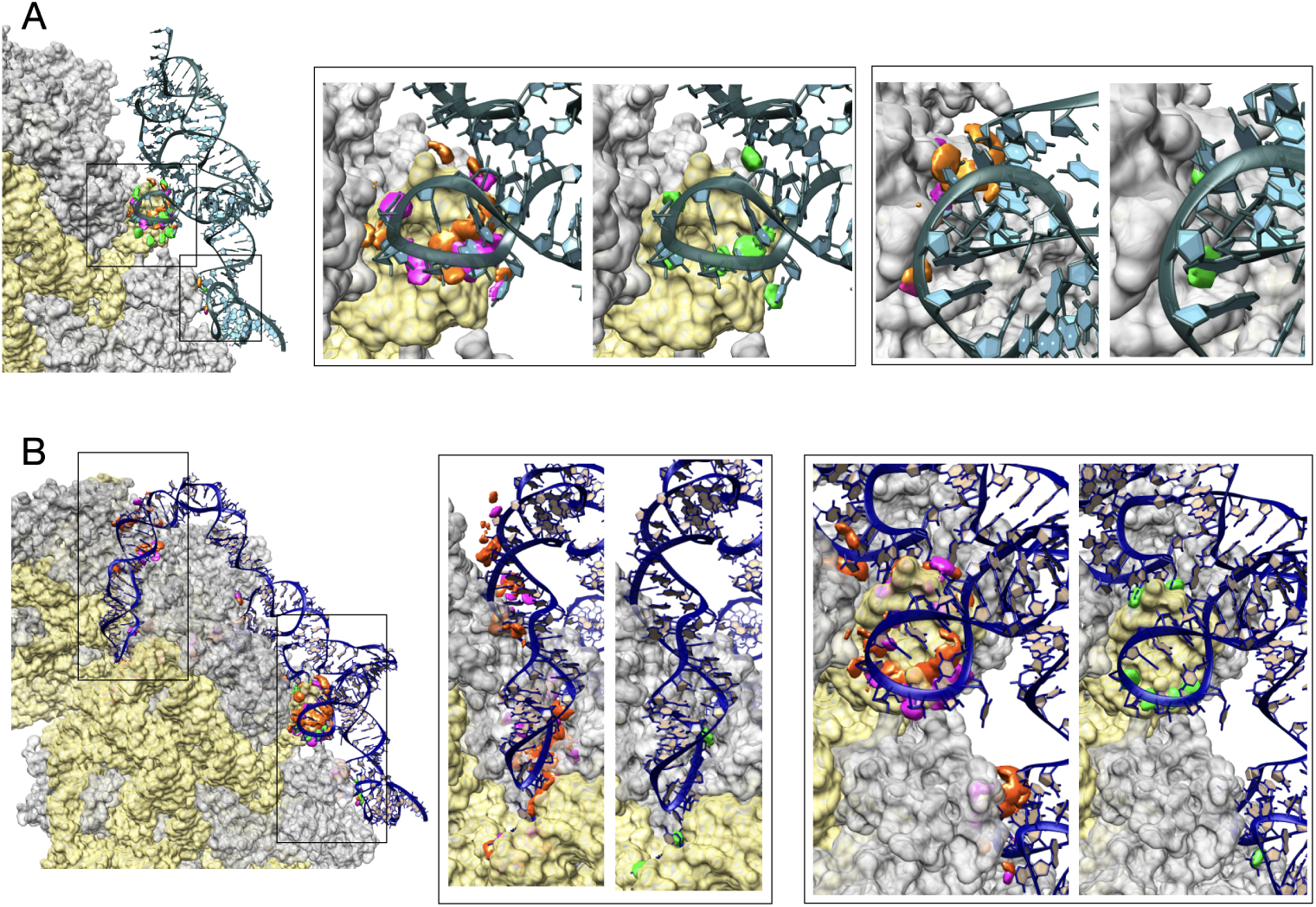
Zoom on the closed (A) and open (B) IRES configuration where we show the results of the stacking (green) and hydrogen bonding (magenta:hba, and orange:hbd) interface fields. The central panel shows the details of the top interface region while the right panel shows the detail of the bottom interface region, shown as black rectangles on the image on the left.

This energy density reflects the spatial distribution of the electrostatic potential. Overall, the electrostatics appear to be generally unfavorable, though the open conformation demonstrates a more favorable interaction profile than the closed conformation. Figure 12 focuses on stacking interactions and HBs, where the isovalue should be interpreted in a more binary fashion, indicating the presence or absence of interactions. For the closed conformation, a couple of small anchor sites are observed where stacking and hydrogen bonding interactions are concentrated. In contrast, the open conformation reveals an additional large patch of interactions (located in the top left quadrant in the figure), prominently involving HB interactions between the ribosome and the head region of the IRES, thereby highlighting the structural differences in interaction networks between these conformations.

## Conclusions

In this work we report our recent development on the calculation and visualization of molecular fields by means of effective potentials based on simple functional forms and parameterized by statistical analysis. Our calculations assume a coarser view than what is usually done for MIFs, where calculations are performed based on detailed atomistic or even quantum force-fields, and don’t use any probes. The calculation and visualization of the stacking field is particularly relevant as the interactions of aromatic rings are key for the binding of RNAs.

We have shown that despite the reduced level of detail of our calculations, our fields are able to capture the core physical features of binding pockets. A benchmark on 20 proteins and RNA systems shows that we obtain a very good agreement between regions of high field values (field hotspots) and the actual placement of known ligands. This agreement is very promising as our tool provides an inexpensive and fast approach to understand and characterize possible binding modes of a pocket that could help in selecting and optimizing binding partners. Our method, grounded in a physical approach, is built on principles that apply equally to both proteins and RNAs, and could help rationalize the main similarities and differences in their interactions with small ligands which, in turns, could allow to better refine computational tools for binding pocket detection and docking for RNAs.

Our approach produces an output that can be promptly visualized with most commonly used visualization software packages such as Chimera, VMD and UnityMol. For the latter, we provide a script that can also be run in a virtual reality context for in-depth exploration of these fields and their use as guidance in interactive docking of ligands or fragments. Calculations are fast enough that fields for a full single macromolecular structure can be obtained in matter of seconds, allowing to visualize the physical properties of a molecule as a whole. This possibility opens the way to use our fields to rationalize the interactions between macromolecules such as two proteins or a protein and a nucleic acid.

## Authors’ contributions

D.B.M developed the approach for APBS, stacking and hydrophobic fields, wrote the original code, performed benchmarks, developed the UnityMol extension, wrote the manuscript; G.M. performed benchmarks, performed the comparison with pharmacophores, developed and applied interface fields, wrote the manuscript; L.M. performed the comparison with other methods and developed visualization plugins. A.K. developed the approach for HBD and HBA; A.R. developed the interface fields; L.R. wrote, documented and tested the distributed code and corresponding websites; H.S. contributed to code re-writing; M.B. supervised the code re-writing, testing and visualization experiments, designed the VR scenario; A.T. conceived applications and wrote the manuscript; S.P. conceived and supervised the work, wrote the manuscript.

## Data availability

Software, benchmarks and demos are available at: https://gitlab.galaxy.ibpc.fr/rouaud/smiffer

Supplementary videos of UnityMol VR applications are available here: https://unity.mol3d.tech/mov/fields_vr1.mp4 https://unity.mol3d.tech/mov/fields_vr2.mp4

Supplementary videos for the IRES system are available here: IRES closed: https://www.youtube.com/watch?v=d5Qu9G08srg IRES open: https://www.youtube.com/watch?v=h7y8t4I8zIY

## Supporting Information

Section 1: Derivation of stacking parameters.

Section 2: Visualization tests and image generation.

Section 3: Comparison with existing MIF software.

Section 4: Additional figures of all fields for all systems.

Section 5: Code specifications.

