## Supplementary Information for "Statistical Molecular Interaction Fields: A Fast and Informative Tool for Characterizing RNA and Protein Binding Pockets"

#### 1. Stacking potential parameters

Stacking potential parameters $\boldsymbol{\mu}_{ST}$ and $\mathbf{S}$ were determined by a statistical analysis of experimental structures. High resolution (< 2.5 Å) protein, protein/ligand, protein/nucleic and protein/nucleic/ligand structures were downloaded from the Protein Data Bank (PDB) [1], resulting in 86532 structures. From these datasets, 22786 unique ligand IDs were observed. The CIF files for these ligands were downloaded from the PDB.

To detect the aromatic interactions of interest to extract model parameters, we focused on ligand/protein interactions and on nucleic bases/protein interactions. A total of 696676 interactions were sampled from the PDB dataset. While it is easy to identify base/aminoacid interactions as they

*

depend only on the identity of the two partners, identifying aromatic interactions for ligands requires the analysis of each ligand separately. To do so, a first approach consisted of parsing the SMILES string of each ligand keeping only those that contained flags for presence of aromatic bonds (i.e. lowercase letters). A second approach consisted in inspecting the bond topology described in the CIF files and consider a ligand to be candidate for aromaticity if its structure had cycles after discarding tetrahedral atoms. Ligands where the two methods coincided (17823 instances) were considered to have an aromatic group for the next steps. Cases where the methods disagreed (1661 instances) were manually inspected to decide whether to consider them aromatic or not [2].

Before sampling the aromatic interactions, all aromatic groups were simplified into their coarse-grained description determining their center of mass and normal vector. In cases where an aromatic group is made by two adjacent aromatic rings (such as tryptophan or adenine), the center of geometry of the whole group is considered as a single point. Two aromatic groups were considered to interact when the distance of their centers of geometry was between 3 Å and 6.5 Å, following the observations of similar data analysis studies [3]. In this case the values of $r$ and $\alpha$ were recorded. To obtain all aromatic interactions present in a PDB file, an all-to-all pairwise calculation was done between the aromatic groups of the system.

#### 2. Visualization tests

For the experiments in UnityMol, we tested our own implementation of three visualization strategies, one of which was specifically developed for the purpose. Each strategy has its own use case with different conditions: a "cloud representation", solid isosurfaces and mesh isosurfaces. The first approach is to visualize clouds of points that would ideally be denser in regions where the values for the potentials are higher, giving to each vertex of the mesh a color that corresponds to its respective potential value. This method normalizes the values of the potentials to assign appropriate color and opacity levels to the points. These meshes are then handled by a GPU shader program that assigns semi-transparent colors to the triangles according to their neighbouring vertices: more saturated and opaque colors imply higher potential values. Alternatively, the potential grids can be visualized using an isosurface representation at a threshold chosen by the user.

Depending on the molecular system and the potentials to be visualized, a given representation method can be more convenient or ergonomic to use. In some instances, for example, superposing the electrostatic potential with a ligand can be confusing when using only wireframe isosurfaces (Fig. S4A.III), as the wireframe obfuscates the ligand and the wireframe colors get mixed together. The cloud representation can be more useful for differentiating the electrostatic regions and the atoms below them, although it might still generate confusion (Fig. S4A.I). In the example of Fig. S4A an appropriate intermediate option is combining solid and translucid isosurfaces (Fig.

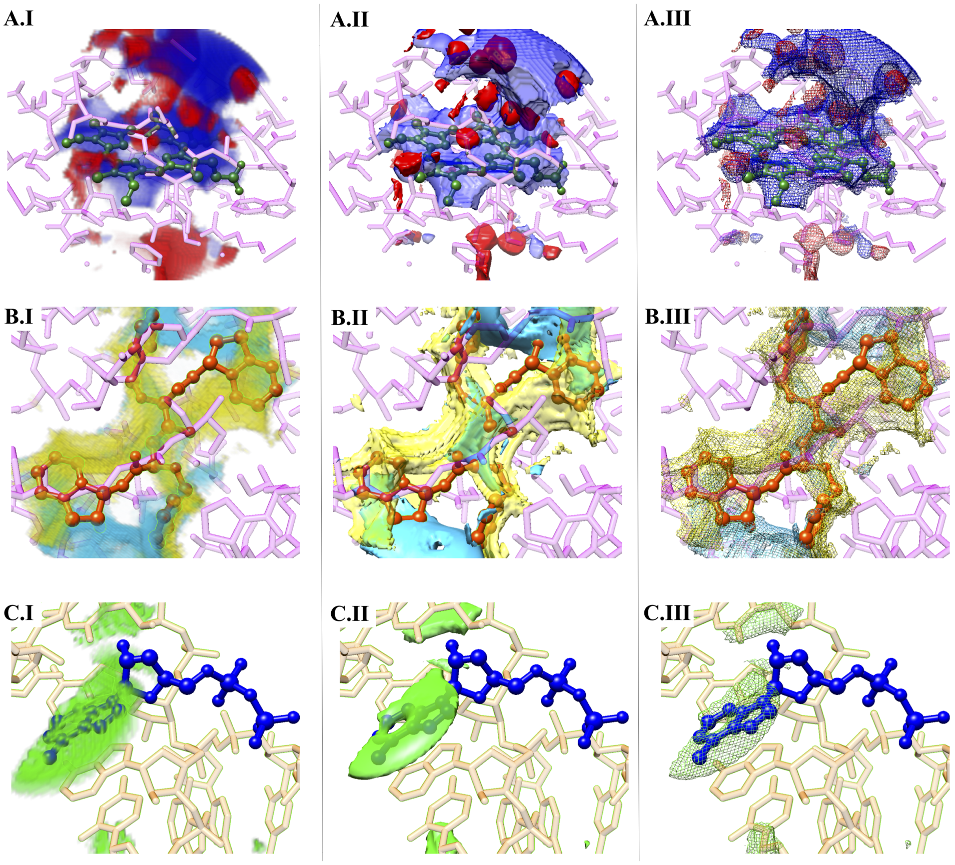


Figure S1: Different representations for visualizing electrostatic potential (A), hydrogen bond potential (B), and stacking potential (C) as I. cloud representation, II. translucid isosurface, III. wireframe isosurface.

S4A.III).

In the case of the stacking potential, comparing the ligand position is even harder when using solid isosurfaces, as it is difficult to distinguish the aromatic rings overlapping with the potential (S4C.III), while translucid and cloud representations are more appropriate for this purpose (S4C.I and (S4C.II). Note however that, if the actual potential shape is of interest and the ligand is disregarded, the solid (not shown) and wireframe isosurfaces are a better choice. Also note that the cloud representation gives less relevance to low value subregions (which can be considered noise), while the translucid isosurface gives the same homogeneous color to the full surface, regardless of the potential values beneath it.

Even if the hydrogen bond potential has some resemblance with the stacking potential in its calculation, the practicality of its representation differs. The cloud and translucid representations are usually vague and not very intuitive (Fig. S4B.I and S4B.II), while the solid and wireframe representations are clearer (Figures S4B.III). This is due to the fact that these potentials are closer to the actual surface of the target (while the stacking extends towards the pocket center), so the ligand is less obstructed during the visualization. The spherical shapes from this potential are very apparent in the solid isosurfaces.

In the current production version of UnityMol, we do only provide a subset of these tested options, namely the isosurface, which can be represented as solid, transparent or wireframe. The script we provide allows to combine and tune these to the users' liking as demonstrated in the supplementary videos.

##### *2.1. Image generation*

Apart from Unitymol, once the SMIFs have been computed and saved, they can alternatively be loaded in a variety of common visualization software packages such as ChimeraX, PyMOL and VMD. For ChimeraX, dropping down the files in the software is enough to visualize them. A Volume viewer automatically opens to adjust the thresholds of the isovalues and various buttons allow to switch representation, from surface, to mesh to volume. For Pymol, only MRC and DX formats are supported, the isovalues can only be modified by editing the mesh/volume levels, which are by default fixed numbers and do not reflect the real values in the file. It is however possible to manage them by commands, for example: cmd.isomesh("1iqj.hbacceptors$_{mesh}$", "1iqj.hbacceptors", 3.0) to set the value to 3. For VMD, Cmap files are not supported, but MRC and DX format files are. The volumes can be loaded as any other files and then thresholds can be controlled using the Graphics Representation tool where one can edit the Isovalue field, change the aspect, the color, as usual with other VMD files. It is also possible to set the visualization in the TkConsole, with "mol modstyle rep$_{number}$ 0 Isosurface 3" to set the value of the map at index 0 to 3.

To make visualization as simple as possible, we developed Plugins for SMIFs for theses three software packages. These are accessible from the SMIFFer software home page. In the future we plan to develop also plugins that launch the calculations of SMIFs directly inside ChimeraX, VMD and Pymol.

#### 3. Comparison with existing MIF software

We present here an example of MIFs computed with three different software packages: GRID [4], MOE [5] and SiteMap [6], [7], for a protein (1IQJ) and one RNA (5XK9). All these software packages make use of probes and compute MIFs based on atomistic force fields like those used in molecular dynamics simulations and docking.

For GRID, results were obtained by first loading the structures into the software, by computing the structures' features and then running the MIF calculations with the "Compute protein MIFS" button. Only the default probes were used for the computation. The resolution used is 1Å . For MOE, we imported the structures via the "Import structure" button, then we used the "Compute → Surface and maps" to compute all the different maps, with the default settings. For Sitemap, we imported the structures in Maestro using "Import Structures", then we applied "Protein Preparation Workflow", with the default settings. We launched a Sitemap Job with the default settings except for the grids, set to fine at 0.35Å . For all the three software packages, the grids of the available results were exported in CCP4 file format, and then opened in ChimeraX.

Three main differences emerge between results of these three software packages and our SMIFs. First, the use of probes, while preserving many details, renders the analysis of the observed fields rather complex as it can be seen for GRID and for MOE. SiteMap uses a postprocessing to recover an information similar to what we propose with our fields. Second, the

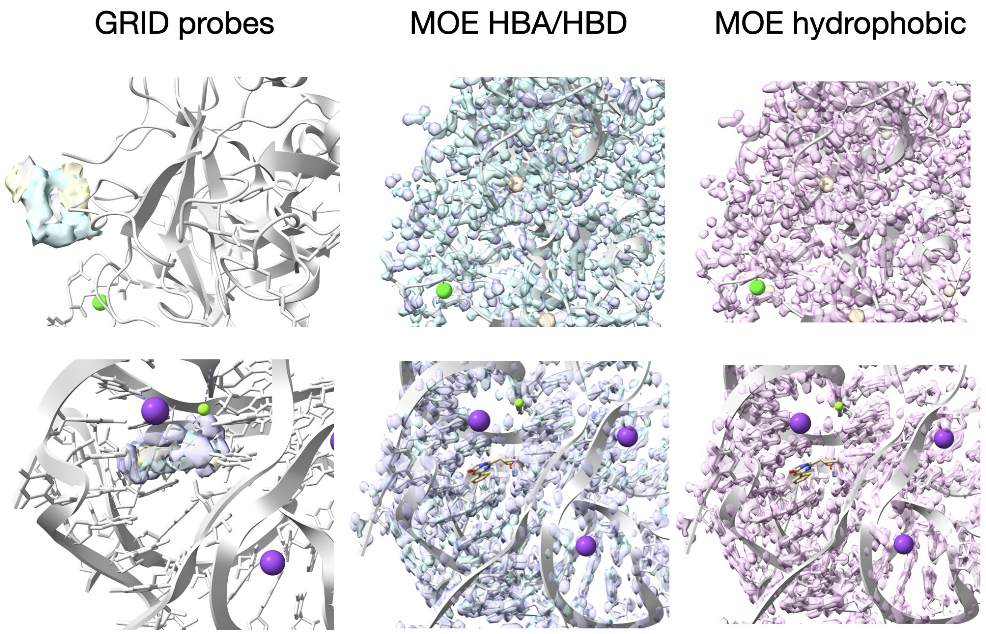


Figure S2: MIFs computed with GRID (probes N, N and O) and MOE for a protein pocket (top) and for an RNA pocket (bottom). MOE automatically computed the MIFs on the whole system.

resolution of these software packages is lower than what we set for SMIFs. Typically MIFs are computed with 1Å resolution, while we compute with 0.25Å resolution. Third, computational times are higher. For the two chosen systems, computation of fields inside the pocket take 11 to 16 seconds with SiteMap, around 20 seconds with MOE, and close to one minute for GRID. It is worth pointing out that all these software packages operate under paying license and are therefore not so easily accessible to the academic community.

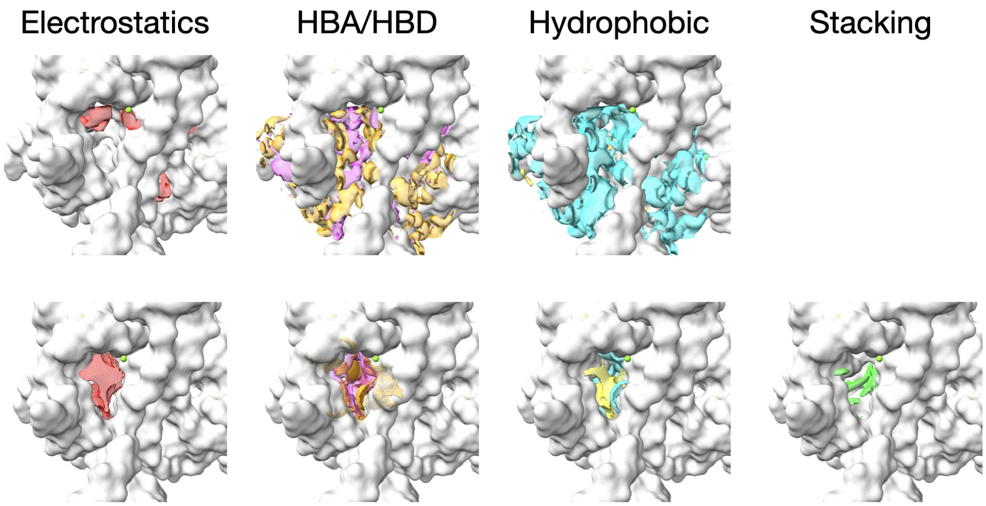


Figure S3: Top: MIFs computed with SiteMap for a protein pocket; Bottom: SMIFs computed with SMIFER for the same pocket.

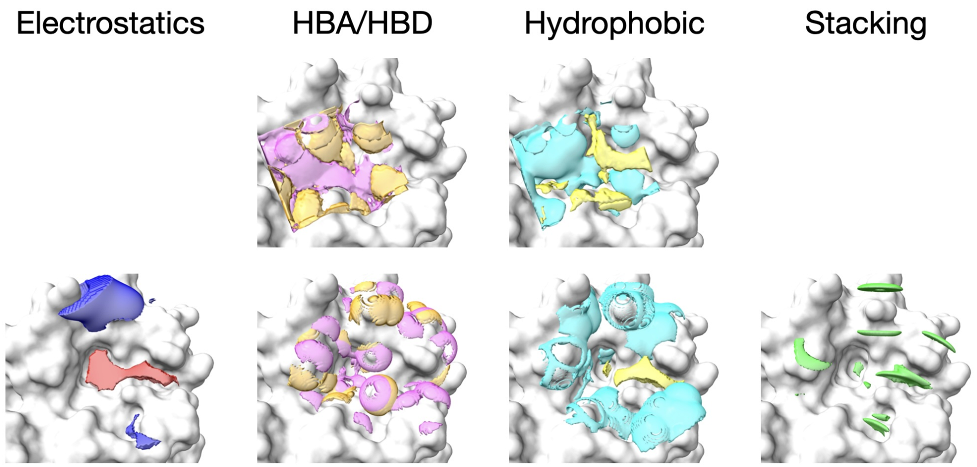


Figure S4: Top: MIFs computed with SiteMap for a protein pocket; Bottom: SMIFs computed with SMIFER for the same pocket.

#### 4. All fields

We present here the SMIFs for all the 20 systems we have analyzed. In each figure color-coding is the following: red → negative electrostatics, blue → positive electrostatics, magenta → HBA, orange → HBD, cyan → hydrophilic, yellow → hydrophobic, green → stacking.

1BG0

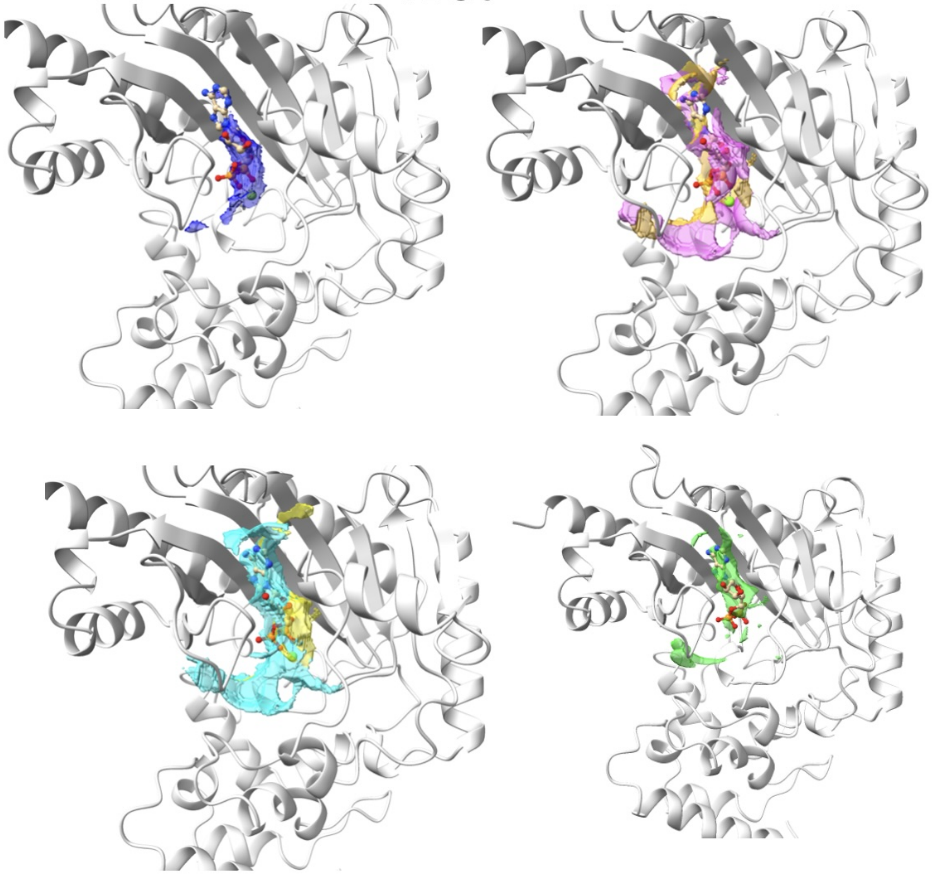

1EBY

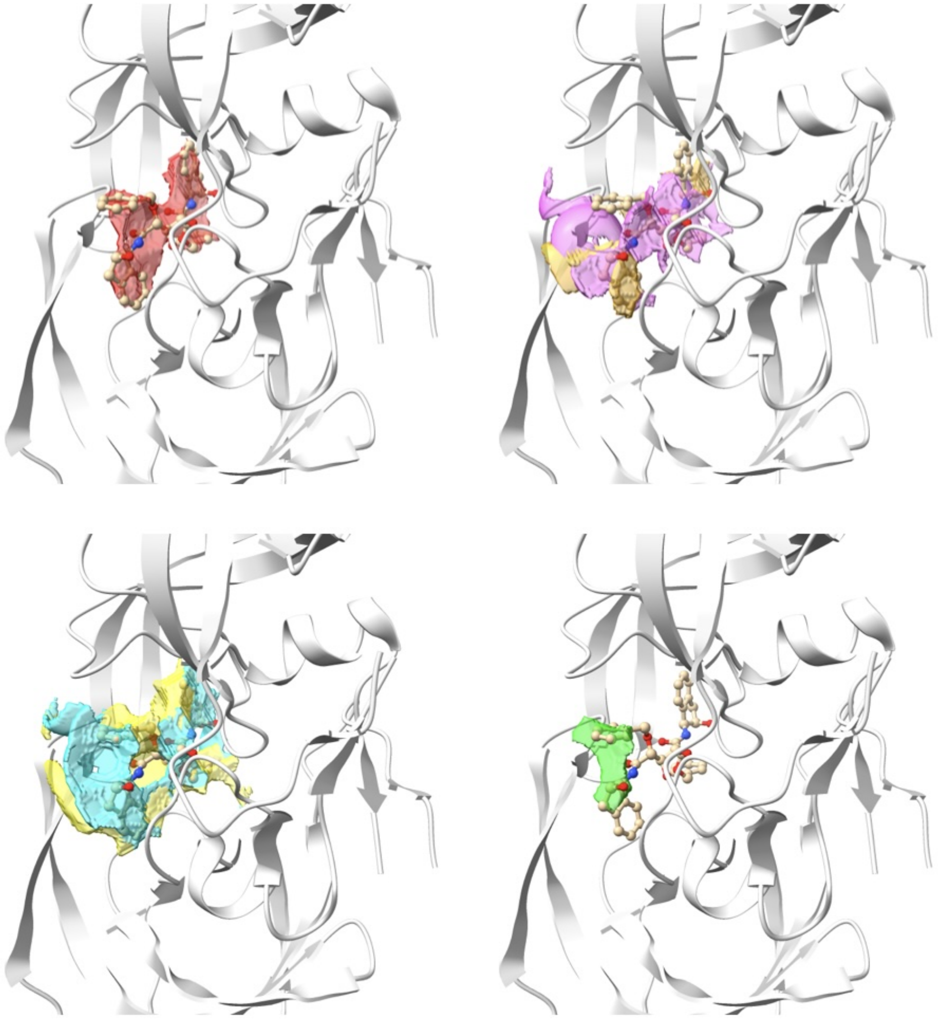

1EHE

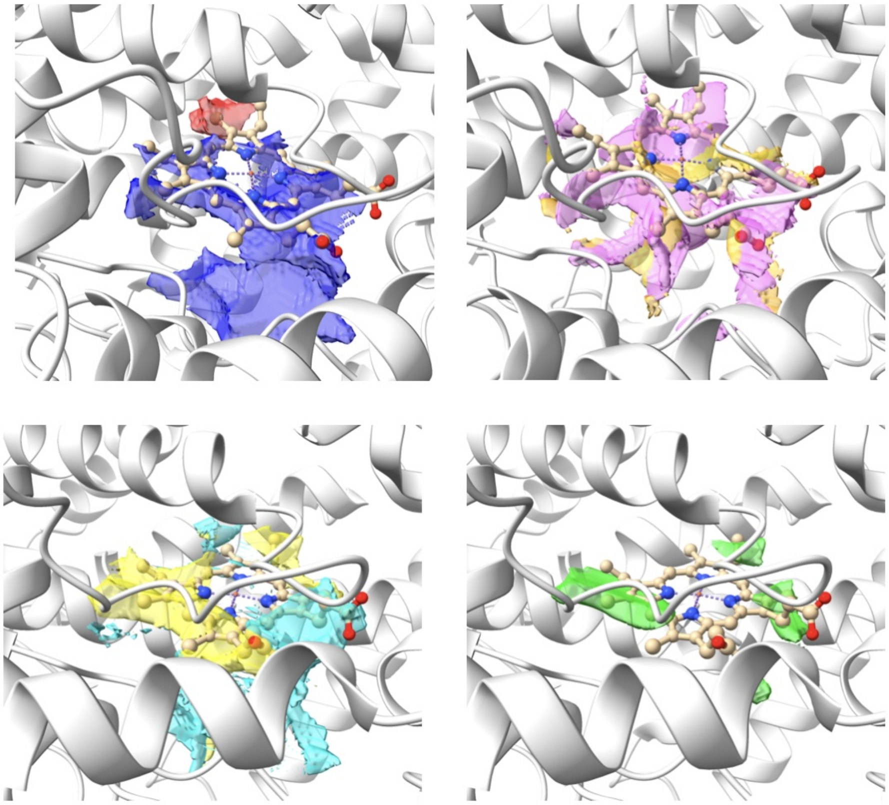

## 1H7I

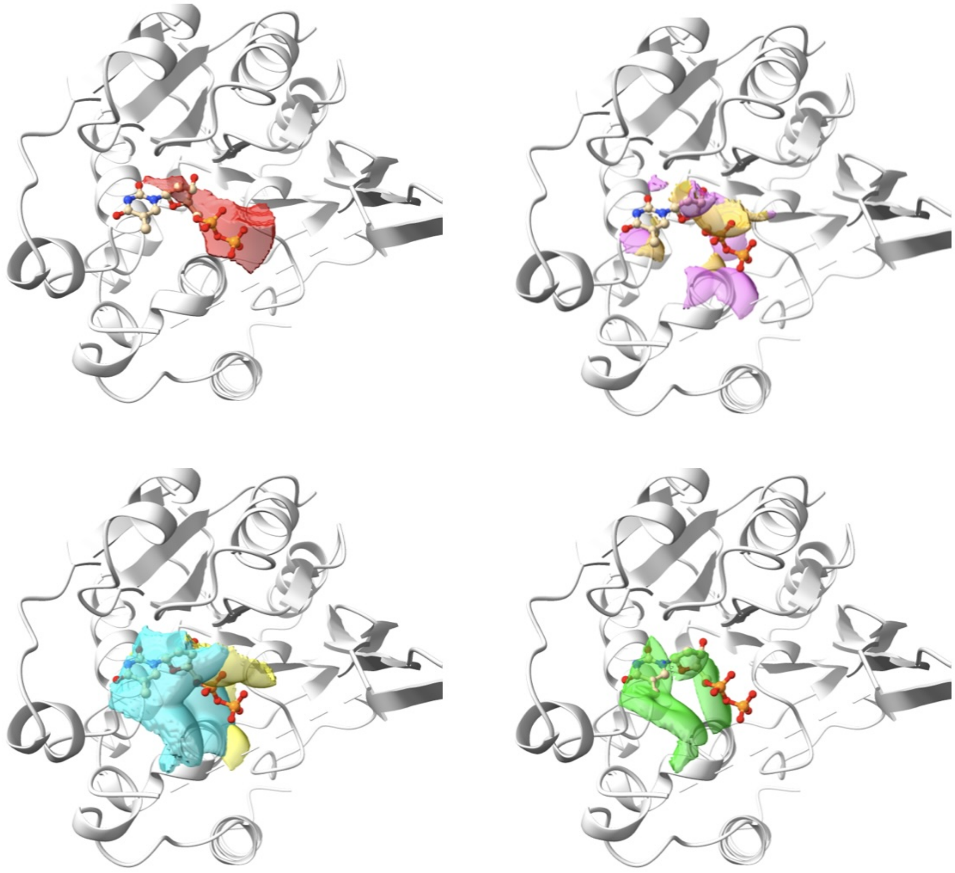

### 1IQJ

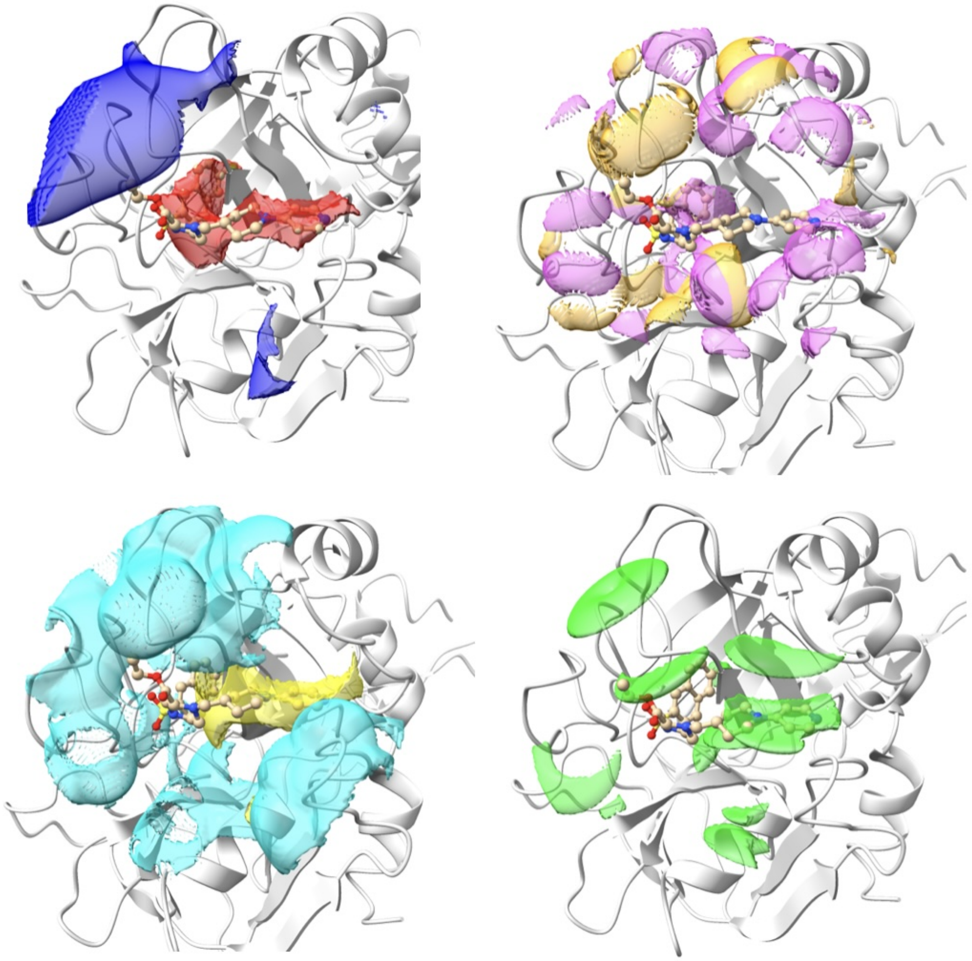

1OFZ

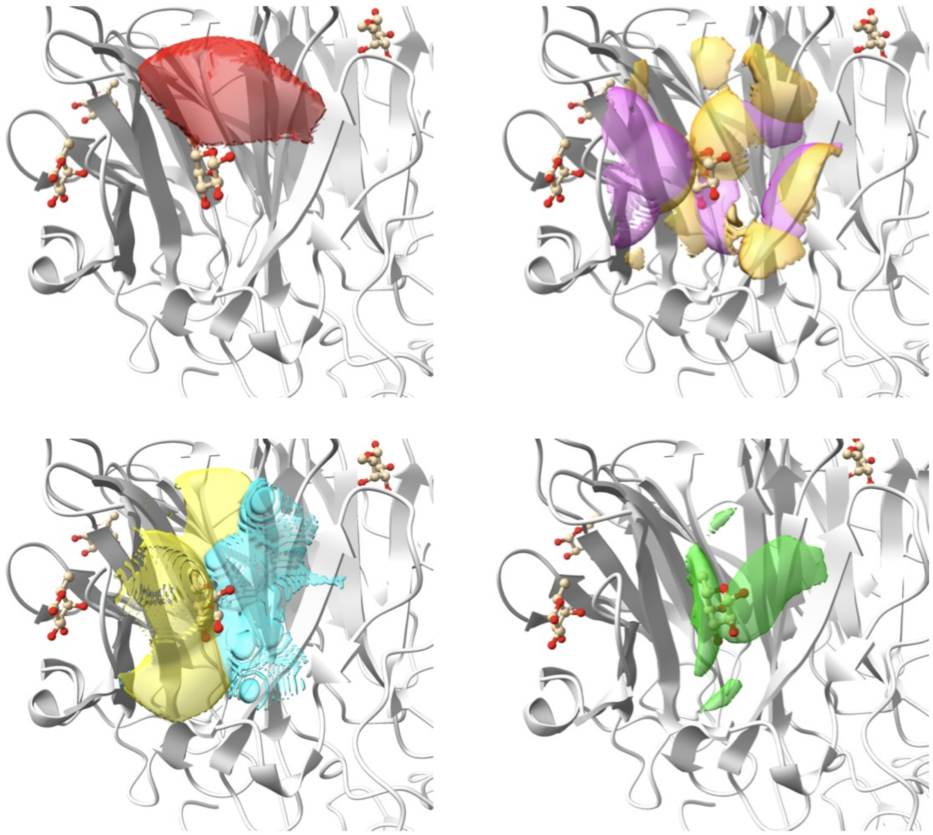

3DD0

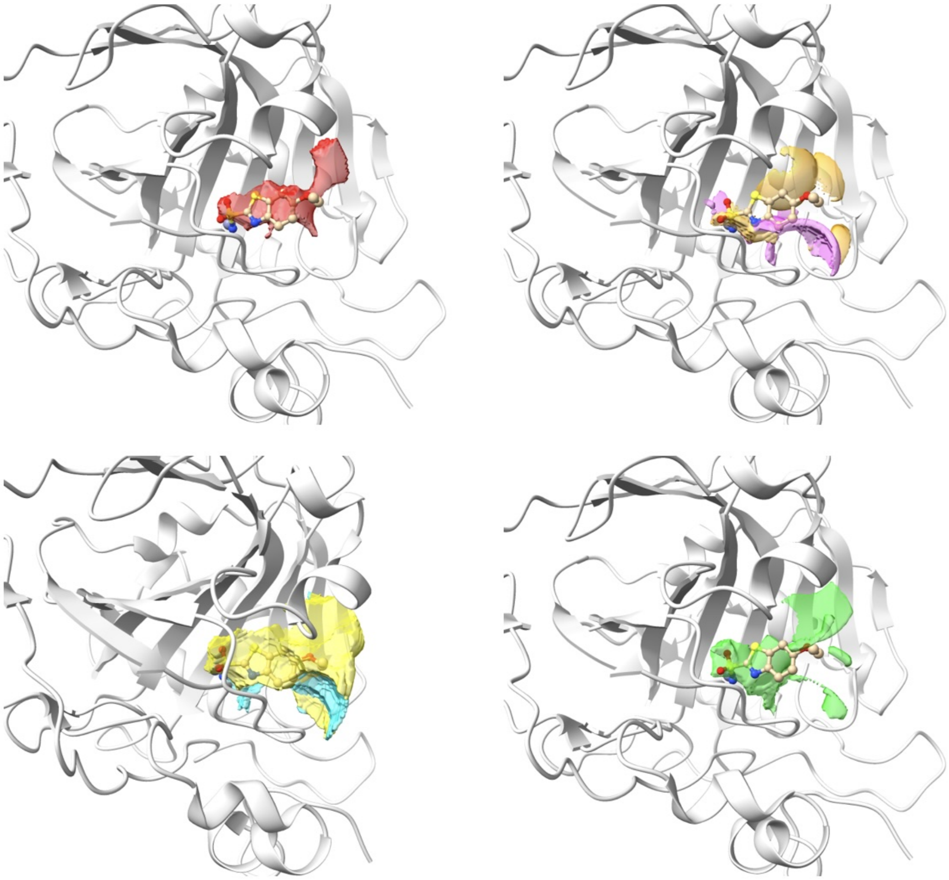

3EE4

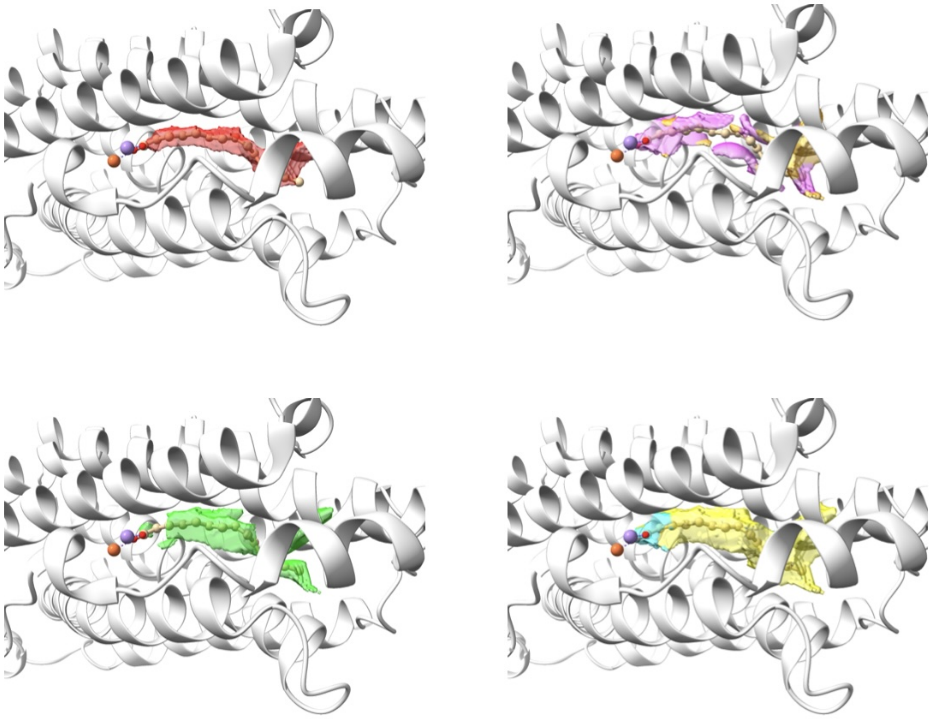

5M9W

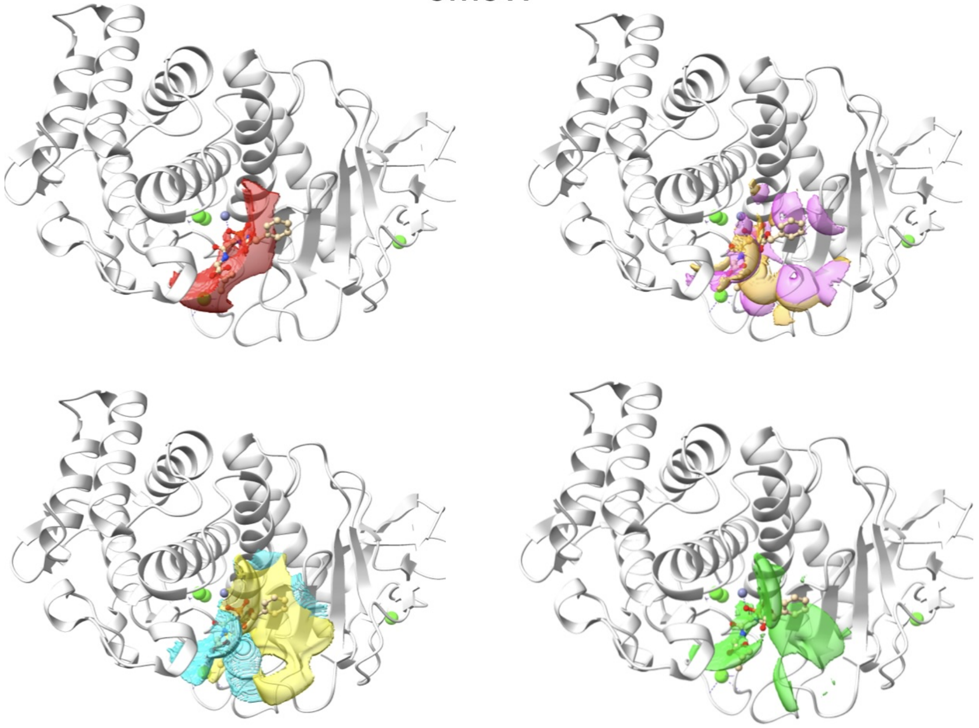

## 6A9A

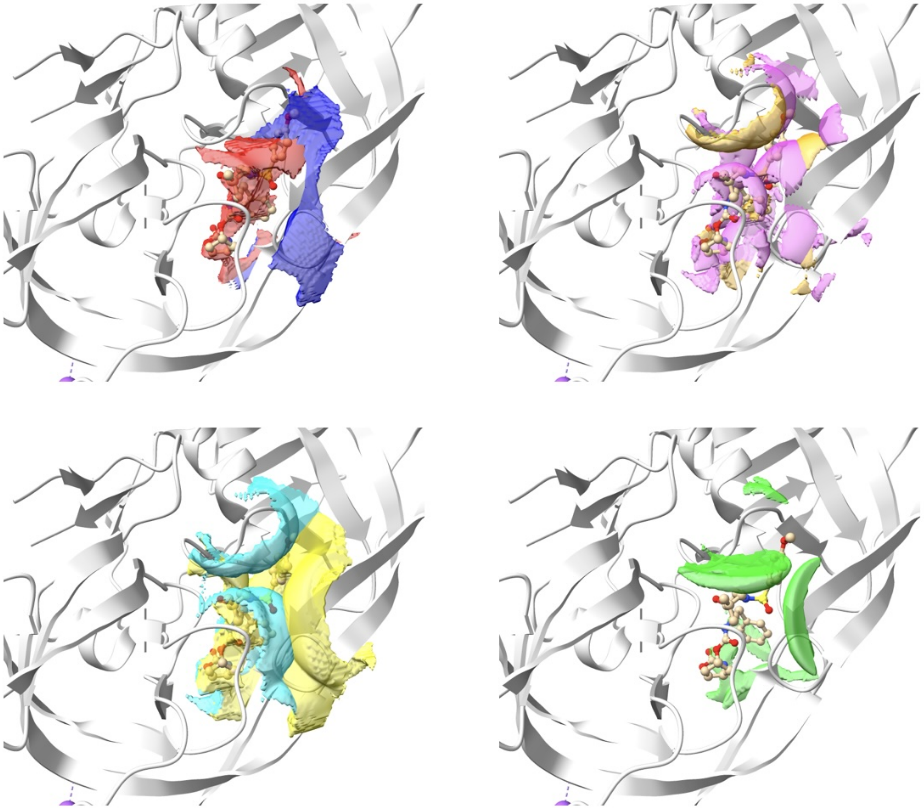

1AKX

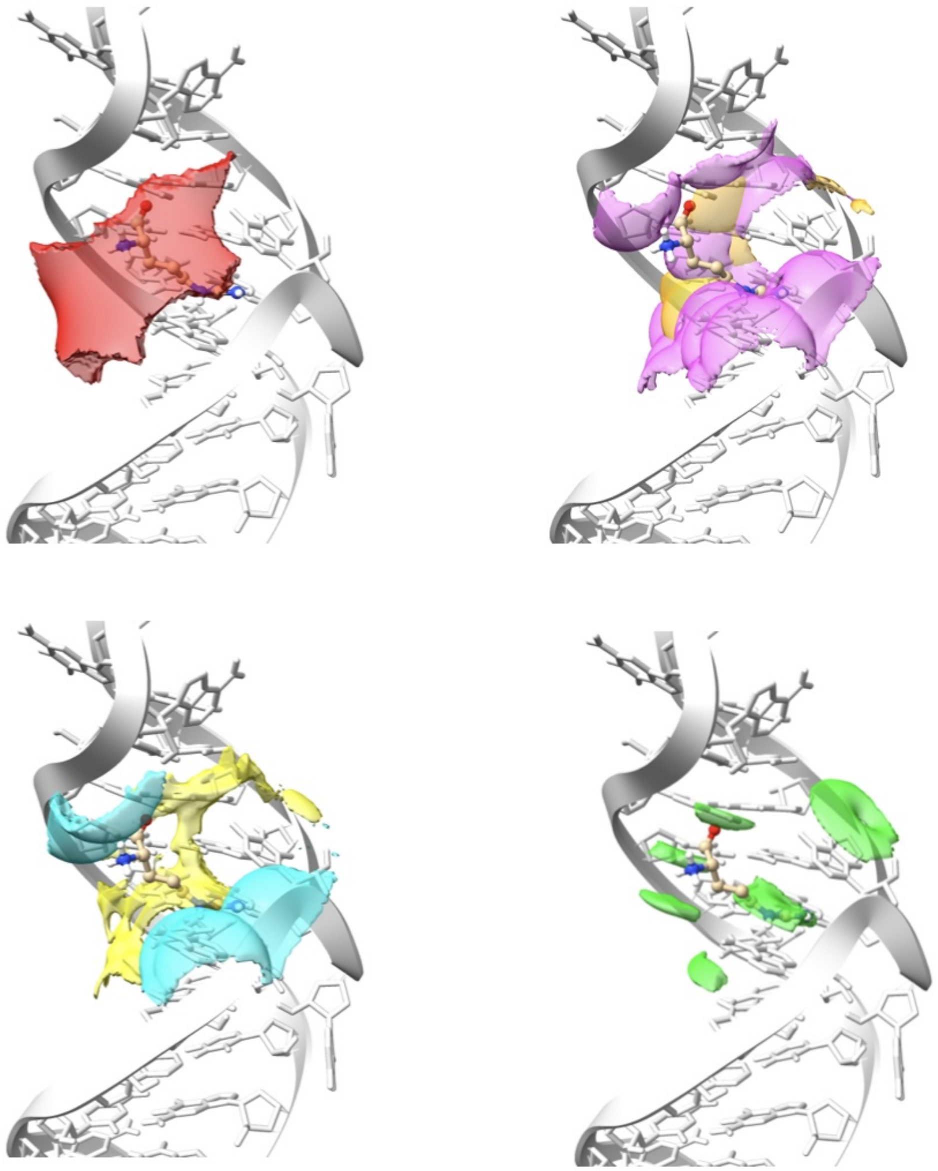

1I9V

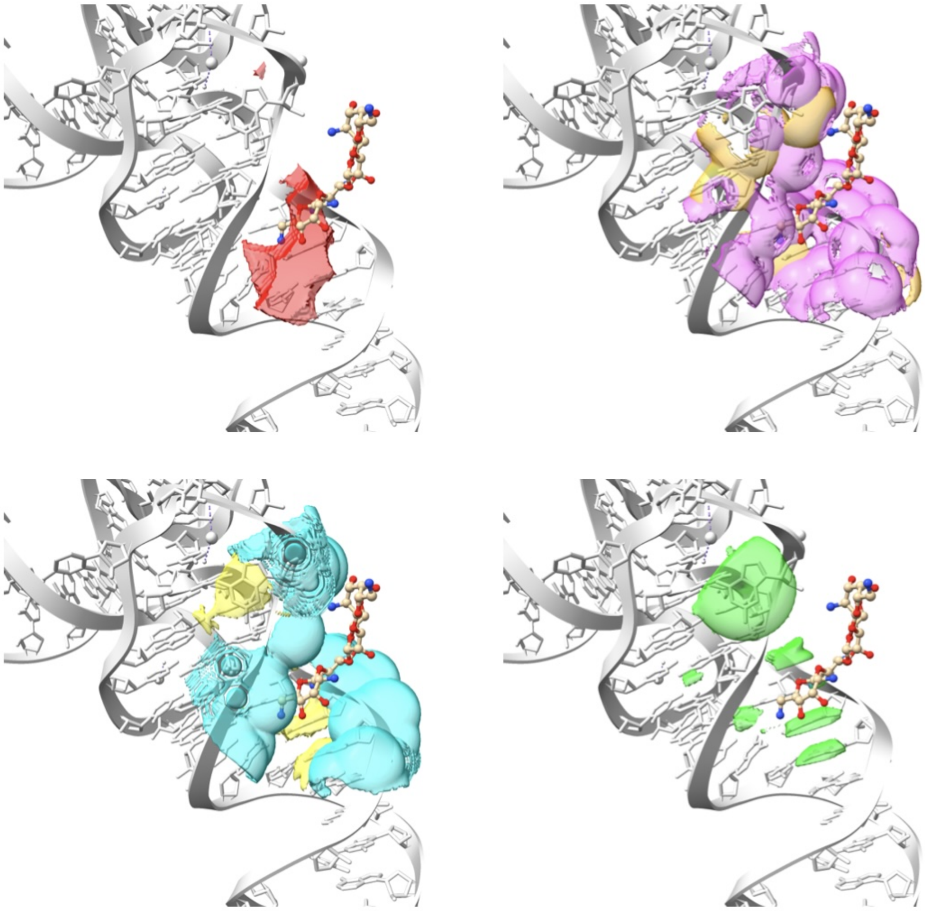

2ESJ

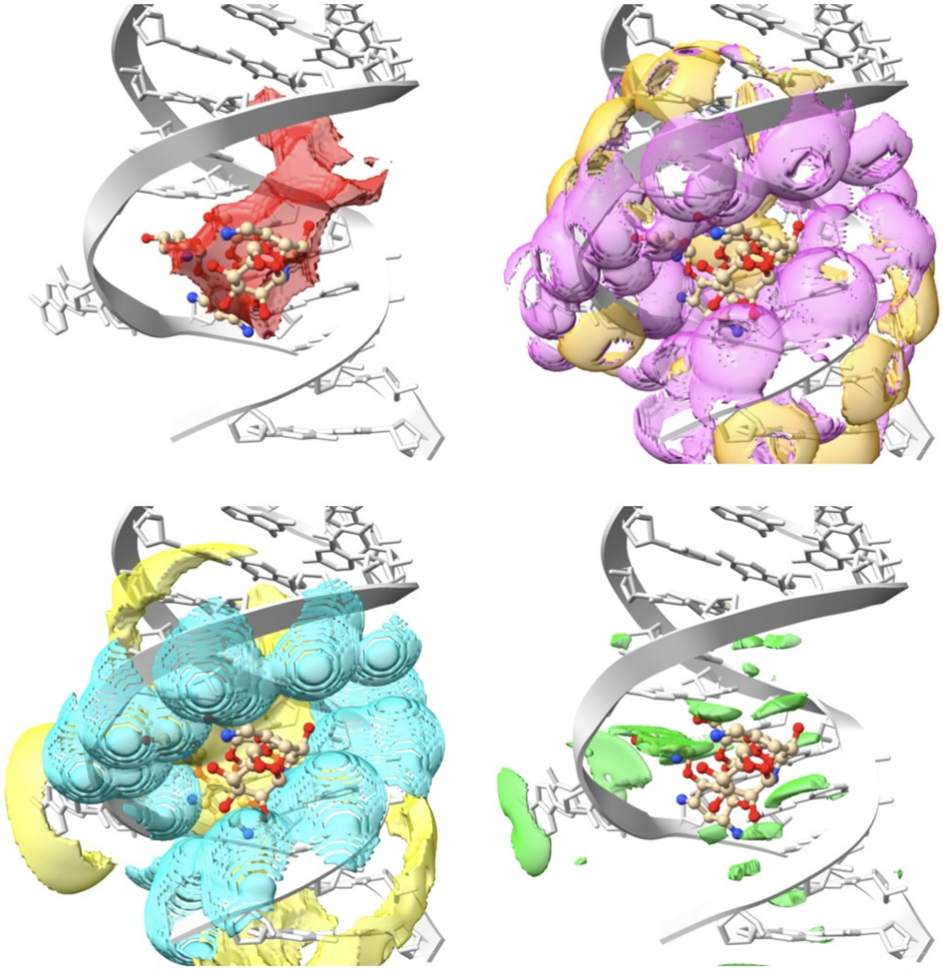

## 4F8U

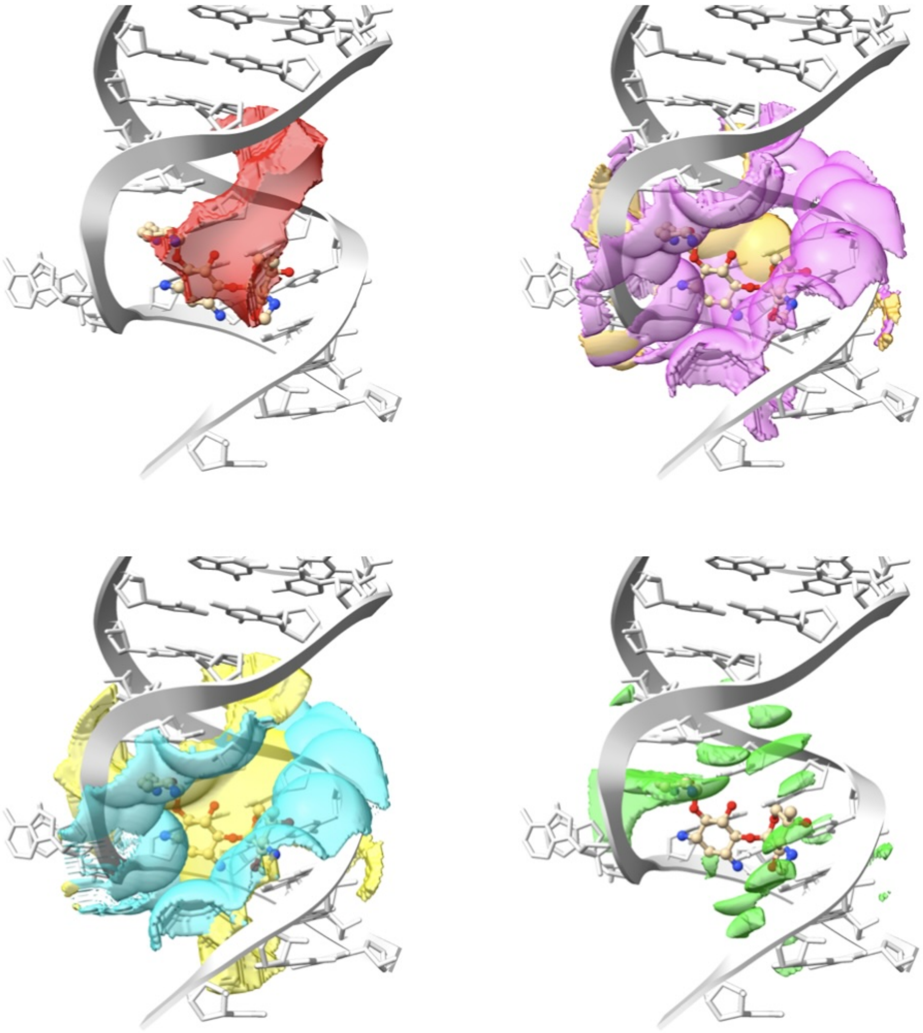

#### 5BJO

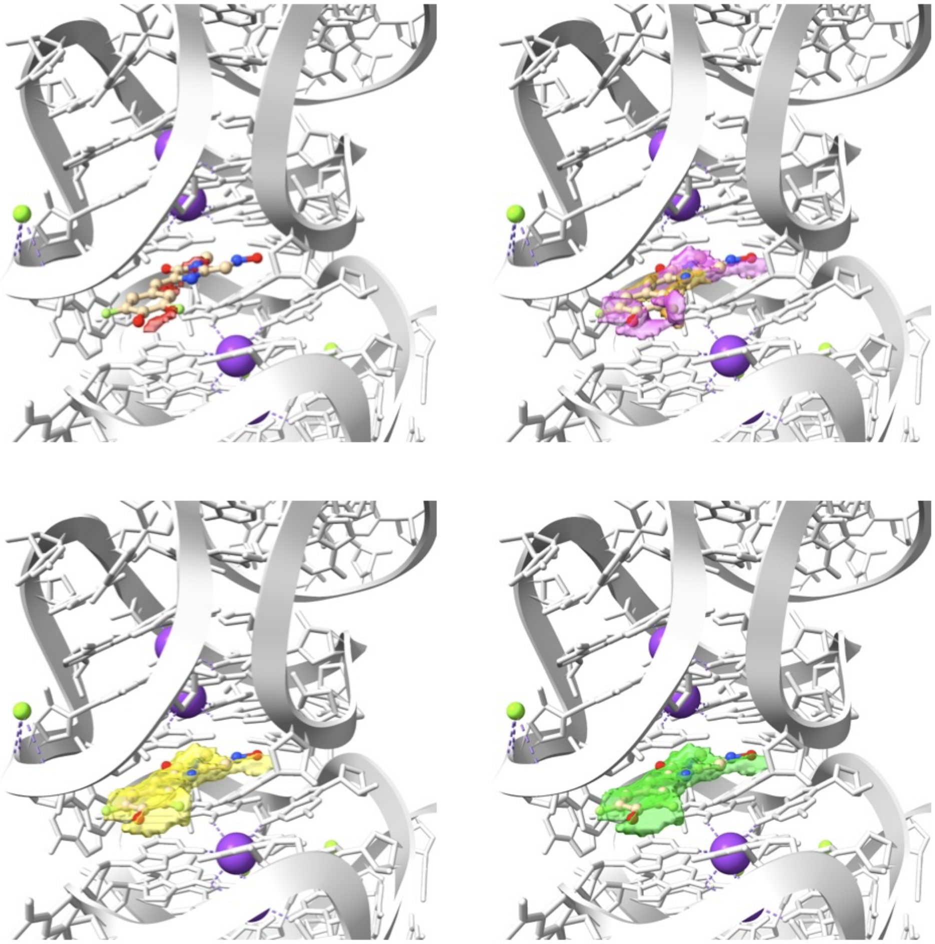

5KX9

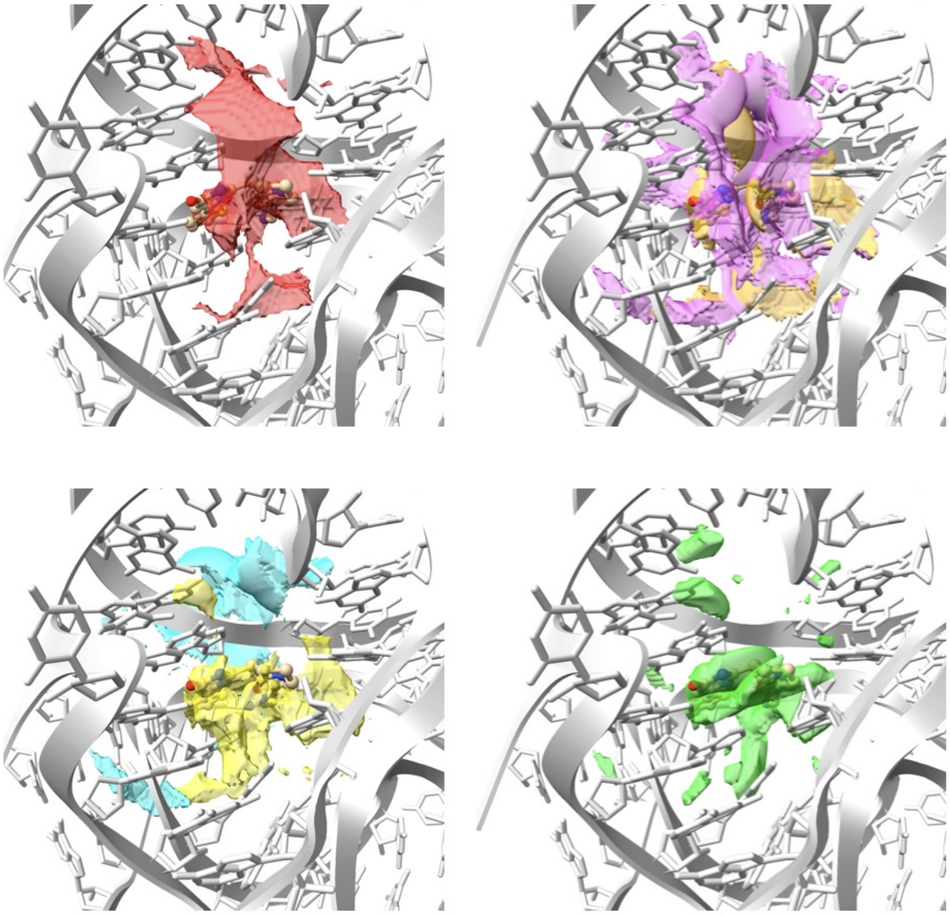

6TF3

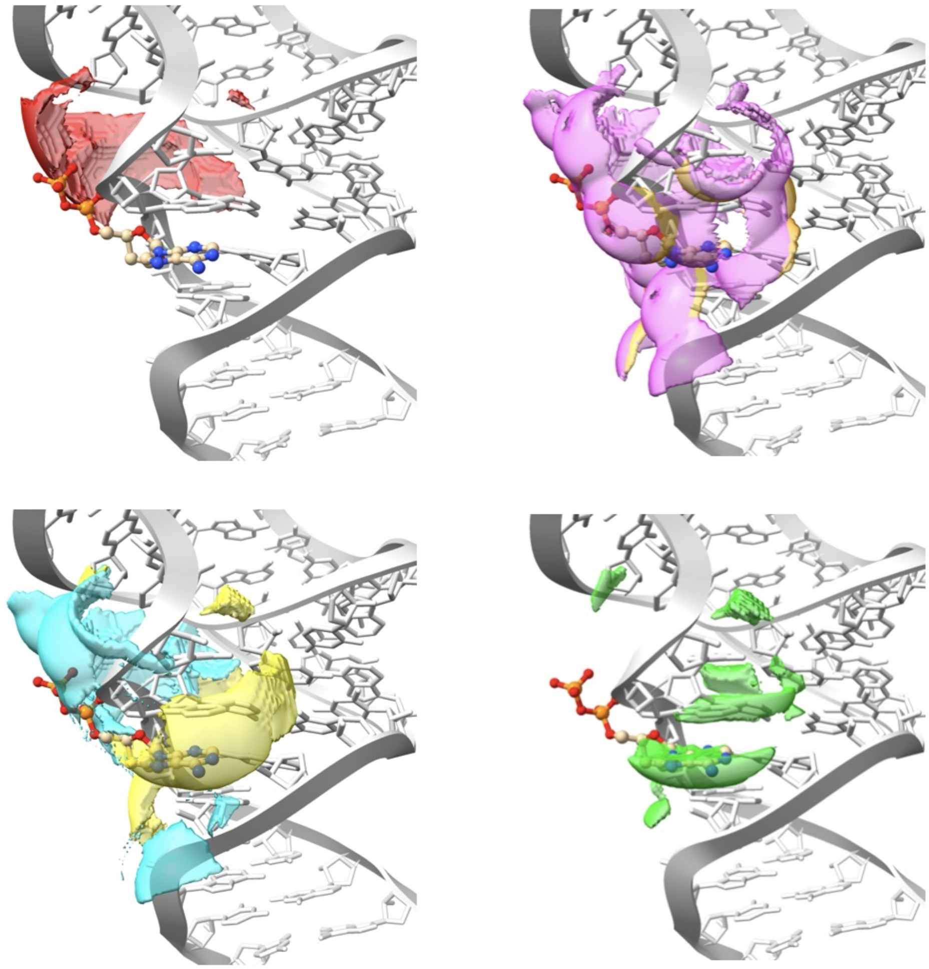

### 7OAX-1

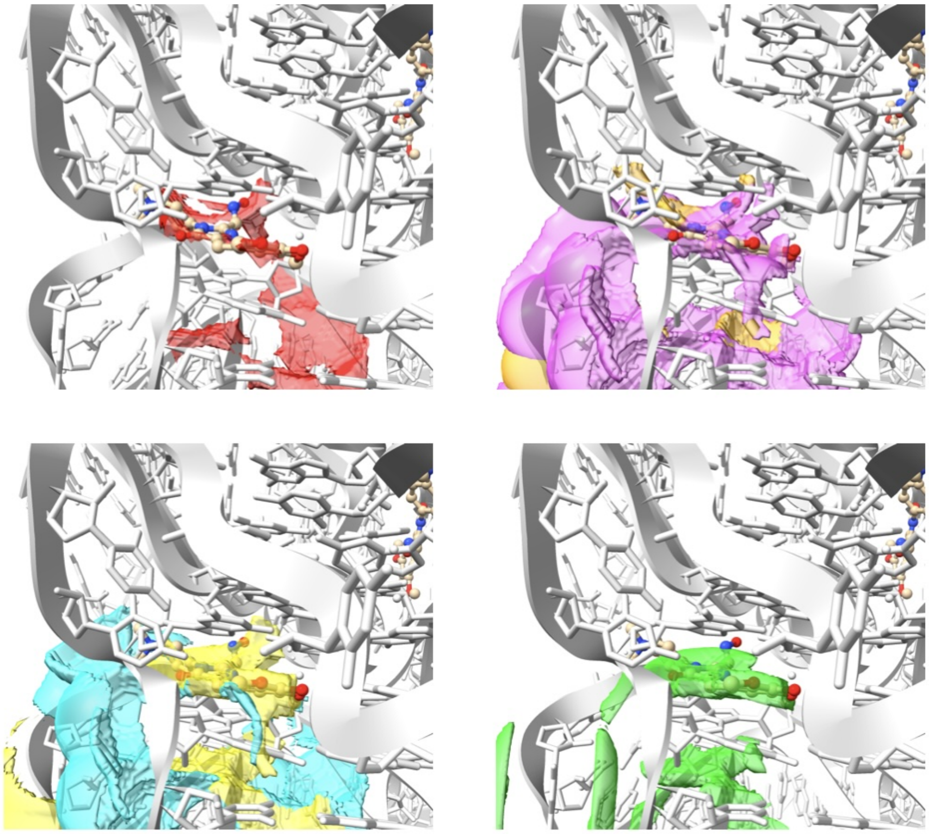

#### 7OAX-2

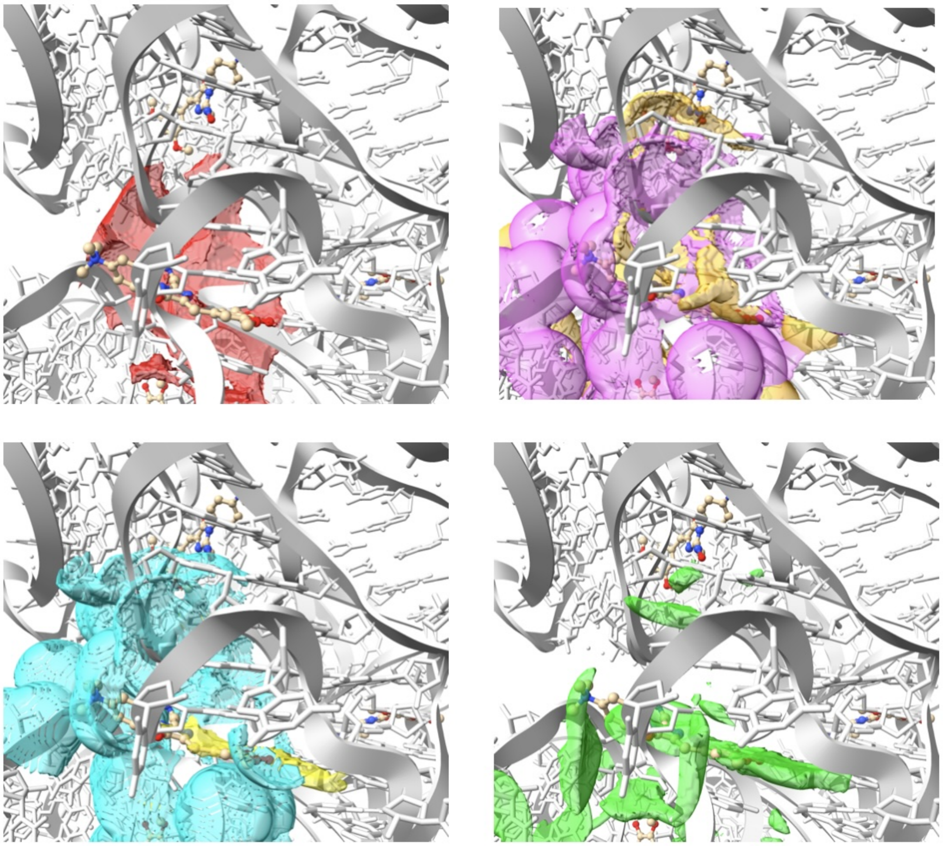

### 8EYV

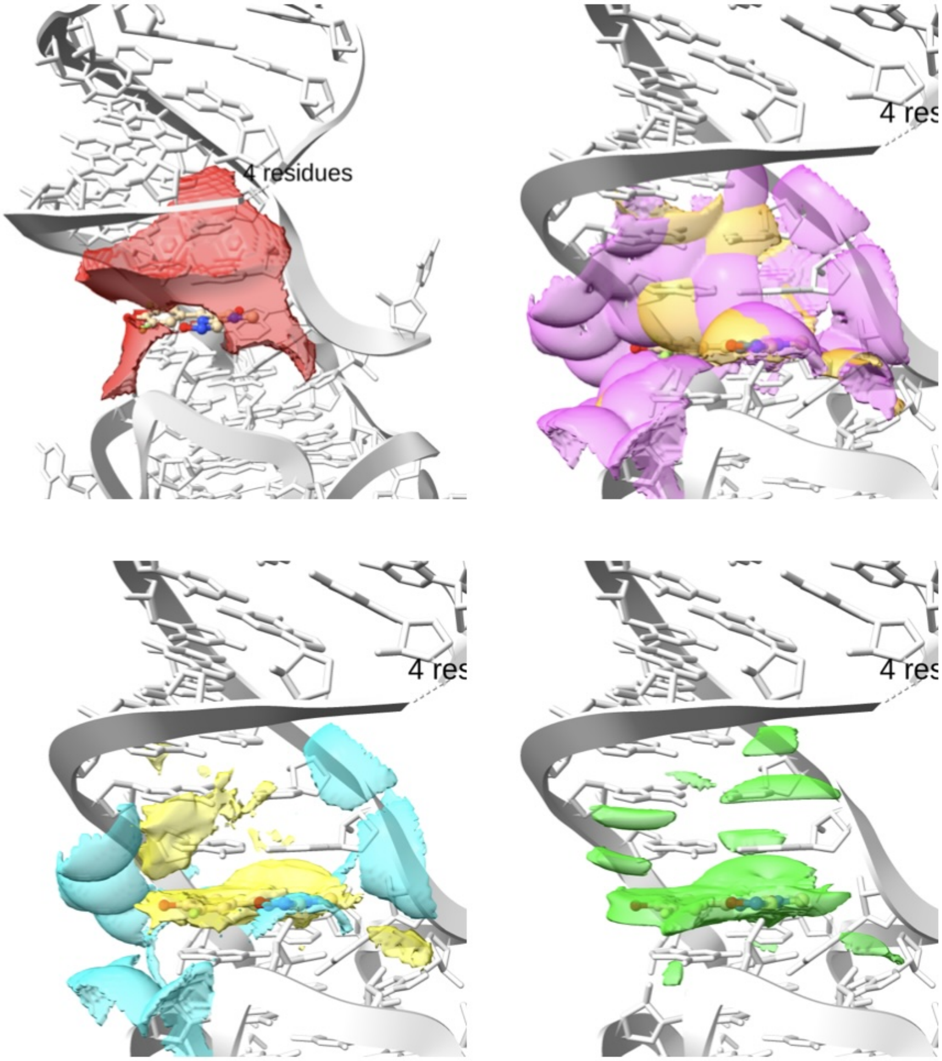

#### 5. Code details

The code can be freely downloaded from gitlab. Here the users find the link to the documentation of the code, as well as a guide on how to install and use the software. A different repository is dedicated to reproduce the benchmarks, together with a report, in *.csv* format, on how much time it takes to calculate and save the fields.

##### *5.1. Code dependencies*

The code is done in python[8] version $\geq$ `3.5`. It relies on the following modules: MDAnalysis[9, 10] version `2.6.1`, used to handle the input PDB; NumPy[11] version `1.26.0`; SciPy[12] version `1.11.4`]; MRCfile[13] version `1.5.3`.

[9] Richard J. Gowers, Max Linke, Jonathan Barnoud, Tyler J. E. Reddy, Manuel N. Melo, Sean L. Seyler, Jan Domański, David L. Dotson, Sébastien Buchoux, Ian M. Kenney, Oliver Beckstein, MDAnalysis:

A Python Package for the Rapid Analysis of Molecular Dynamics Simulations, in: Sebastian Benthall, Scott Rostrup (Eds.), Proceedings of the 15th Python in Science Conference, 2016, pp. 98 – 105. doi:10.25080/Majora-629e541a-00e.

[10] N. Michaud-Agrawal, E. J. Denning, T. B. Woolf, O. Beckstein, Md-analysis: A toolkit for the analysis of molecular dynamics simulations, Journal of Computational Chemistry 32 (10) (2011) 2319–2327. arXiv:https://onlinelibrary.wiley.com/doi/pdf/10.1002/jcc.21787, doi:https://doi.org/10.1002/jcc.21787.
URL https://onlinelibrary.wiley.com/doi/abs/10.1002/jcc.21787

[11] C. R. Harris, K. J. Millman, S. J. van der Walt, R. Gommers, P. Virtanen, D. Cournapeau, E. Wieser, J. Taylor, S. Berg, N. J. Smith, R. Kern, M. Picus, S. Hoyer, M. H. van Kerkwijk, M. Brett, A. Haldane, J. F. del Río, M. Wiebe, P. Peterson, P. Gérard-Marchant, K. Sheppard, T. Reddy, W. Weckesser, H. Abbasi, C. Gohlke, T. E. Oliphant, Array programming with NumPy, Nature 585 (7825) (2020) 357–362. doi:10.1038/s41586-020-2649-2.
URL https://doi.org/10.1038/s41586-020-2649-2

[12] P. Virtanen, R. Gommers, T. E. Oliphant, M. Haberland, T. Reddy, D. Cournapeau, E. Burovski, P. Peterson, W. Weckesser, J. Bright, S. J. van der Walt, M. Brett, J. Wilson, K. J. Millman, N. Mayorov, A. R. J.

Nelson, E. Jones, R. Kern, E. Larson, C. J. Carey, İ. Polat, Y. Feng, E. W. Moore, J. VanderPlas, D. Laxalde, J. Perktold, R. Cimrman, I. Henriksen, E. A. Quintero, C. R. Harris, A. M. Archibald, A. H. Ribeiro, F. Pedregosa, P. van Mulbregt, SciPy 1.0 Contributors, SciPy 1.0: Fundamental Algorithms for Scientific Computing in Python, Nature Methods 17 (2020) 261–272. doi:10.1038/s41592-019-0686-2.

[13] T. Burnley, C. M. Palmer, M. Winn, Recent developments in the ccp-em software suite, Acta Crystallographica Section D: Structural Biology 73 (6) (2017) 469–477.
